# Probe Sensitivity to Cortical versus Intracellular Cytoskeletal Network Stiffness

**DOI:** 10.1101/445320

**Authors:** Amir Vahabakashi, Chan Young Park, Kristin Perkumas, Zhiguo Zhang, Emily K. Deurloo, Huayin Wu, David A. Weitz, W Daniel Stamer, Robert D. Goldman, Jeffrey J. Fredberg, Mark Johnson

## Abstract

In development, wound healing, and pathology, cell biomechanical properties are increasingly recognized as being of central importance. To measure these properties, experimental probes of various types have been developed, but how each probe reflects the properties of heterogeneous cell regions has remained obscure. To better understand differences attributable to the probe technology, as well as to define the relative sensitivity of each probe to different cellular structures, here we took a comprehensive approach. We studied two cell types --Schlemm’s canal (SC) endothelial cells and mouse embryonic fibroblasts (MEFs) – using four different probe technologies: 1) atomic force microscopy (AFM) with sharp-tip; 2) AFM with round-tip; 3) optical magnetic twisting cytometry (OMTC); and 4) traction microscopy (TM). Perturbation of SC cells with dexamethasone treatment, a-actinin overexpression, or Rho-A overexpression caused increases in traction reported by TM and stiffness reported by sharp-tip AFM, as compared to corresponding controls. By contrast, under these same experimental conditions, stiffness reported by round-tip AFM and by OMTC indicated little change. Knock out (KO) of vimentin in MEFs caused a diminution of traction reported by TM, as well as stiffness reported by sharp-tip and round-tip AFM. However, stiffness reported by OMTC in vimentin KO MEFs was greater than in wild-type. Finite element analysis demonstrated that this paradoxical OMTC result in vimentin KO MEFs could be attributed to reduced cell thickness. Our results also suggest that vimentin contributes not only to intracellular network stiffness but also cortex stiffness. Taken together, this evidence suggests that AFM sharp-tip and TM emphasize properties of the actin-rich shell of the cell whereas round-tip AFM and OMTC emphasize those of the non-cortical intracellular network.

## INTRODUCTION

Biomechanical properties of cells influence a variety of basic cellular behaviors including migration, proliferation, contraction, elongation, adhesion, cytokinesis, and apoptosis (1–3). These behaviors in turn impact tissue morphogenesis, cancer metastasis, bronchospasm, vasospasm, systemic hypertension, glaucoma and atherosclerosis (4–10). Early studies of cellular biomechanics considered the cell to be either a homogeneous structure or a fluid-like cytoplasm surrounded by a cortical elastic shell. It is now clear, however, that the mechanical behavior of a cell is determined by the structure of the cortical, intracellular (non-cortical) cytoskeletal, and nuclear networks (11, 12), their distribution in space, and the complex rheology of the cytoplasm and cytosol (13). But how these factors are reflected by different experimental probes remains unclear.

We examine here the cytoskeletal mechanics of endothelial cells of the inner wall of Schlemm’s canal (SC cells), with possible relevance to glaucoma (14, 15). To expand the generality of the results, and to focus on the role of vimentin, we also consider mouse embryonic fibroblasts (MEFs). Experiments employed atomic force microscopy (AFM) with both sharp and round tips, optical magnetic twisting cytometry (OMTC) and traction microscopy (TM). Each of these probes provide complementary characterization of the material properties of these cells with AFM tips measuring resistance to indentation (12), OMTC measuring the resistance of the cells to applied torque (16), and TM characterizing the active tensile component in the cells (17). These are surface probes that would be expected to be influenced by the stiffness of the F-actin dense cortical region of the cell, which we refer to here as the cortex, and by the intracellular cytoskeleton, but not by the much softer cytoplasm/cytosol (11, 18).

We found that for all manipulations to both cell types, sharp-tip AFM probes and TM gave similar characterization of cell biomechanical behavior and were sensitive primarily to the cortical region of the cell. In contrast, larger round-tip AFM probes and OMTC were sensitive to both the cortical and intracellular network, but primarily the latter. Finally, the absence of vimentin caused a reduction in stiffness of both the cortical and intracellular networks and in cellular traction.

## MATERIALS AND METHODS

### Cell Isolation and Culture

SC cells were isolated and cultured from post-mortem enucleated human eyes within 36 h of death with enucleation occurring less than 6 h after death (15). Eyes with history of trabeculoplasty, anterior segment surgery or any ocular disease (except cataract) were excluded. SC cells between passages 3-5 were used for all experiments. SC cells were characterized by established procedures (expression of vascular endothelial cadherin, net transendothelial electrical resistance of 10 ohms · cm^2^ or greater, and lack of myocilin induction by dexamethasone (15)). SC cells were cultured in DMEM/Low glucose (Life Technologies, Grand Island, NY) with 10% fetal bovine serum (Atlanta Biologicals, Norcross, GA) and 1% penicillin/streptomycin (Life Technologies, Grand Island, NY).

Wild type (WT) and vimentin knock-out MEFs were originally extracted from mouse embryo and immortalized as discussed previously (19). MEFs were cultured in DMEM/High glucose (Life Technologies, Grand Island, NY) supplemented with 10% fetal calf serum (FCS) (Atlanta Biologicals, Norcross, GA), 5 mM nonessential amino acids (NEAA) (Life Technologies, Grand Island, NY), and 1% penicillin/streptomycin (Life Technologies, Grand Island, NY). MEFs were either sparsely seeded or confluent when tested.

All cell cultures were maintained in a humidified incubator at 5% CO2 and 37 °C, and media was changed every other day. Cells were passaged when they were around 80% confluent. For passaging, cells were washed with PBS (Life Technologies, Grand Island, NY) to remove the serum, then treated with trypsin-EDTA 0.25% (Thermo Fischer Scientific, Grand Island, NY), and passaged at a ratio of 1:3.

Cell cultures were examined under various conditions of confluency: superconfluent, confluent or sparsely confluent.

### Dexamethasone Studies

Stock solutions of dexamethasone (0.01, 0.1, and 1 mM; Sigma Aldrich, Milwaukee, WI) were prepared in ethanol as the vehicle and then diluted with culture medium to final concentrations of 0.01, 0.1 and 1 μM dexamethasone. Control studies used the same ethanol concentration (1%) as used in the dexamethasone solutions. Two normal SC cell strains (SC71, SC76) were treated with dexamethasone or control solutions for seven days as it takes this long to get a reliable dexamethasone response (20). Media was replaced every other day. Cells were superconfluent at the end of the treatment.

### Overexpression of α-Actinin and RhoA Expression

Four normal (SC69, SC71, SC73, SC76) cell strains were transduced with (i) an adenovirus encoding a Ubiquitin promoter driving GFP, (ii) a Ubiquitin promoter driving α -actinin with a GFP tag, or (iii) a Ubiquitin promoter driving RhoA with a GFP tag; a fourth group (iv) consisting of non-transduced cells (no virus) was also used. Groups (iii) and (iv) served as controls. Cells were confluent when tested.

### AFM Measurements

All AFM measurements were made with cells in media with 10% fetal bovine serum. AFM measurements were made on a BioScope II with Nanoscope V controller (Bruker, Santa Barbara, CA) coupled to an inverted fluorescent microscope with 10x (NA=0.3), 20x (NA=0.8) objective lens (Carl Zeiss, Thornwood, NY). Sharp tips were pyramidal cantilevers mounted on a triangular cantilever with a length of 200 μm, a nominal tip radius of 20 nm, and a nominal spring constant of 0.02 N/m (Olympus TR400PSA, Asylum Research, Goleta, CA). Round tips were spheres of 10 μm diameter mounted on silicon nitride cantilevers with nominal spring constant of 0.01 N/m (Novascan Technologies, Ames, Iowa). Indentation depths were between 100 and 400 nm that we have previously shown to give values of cellular Young’s modulus (E) that were relatively independent of indentation (12), while also avoiding any substrate effects (21). Measurements were done on regions well away from both nucleus and cell edges. Young’s modulus was determined as described in the Supplementary Information (12). AFM measurements are done with a ramp of 800 nm/sec; we have previously shown that at this speed our measurements are not rate dependent (12), and that with the algorithm used (21), elastic modulus is relatively independent of indentation/load (12). The spring constant was calibrated before each experiment using the thermal fluctuations function of the Nanoscope that measures the motion of cantilever in response to thermal noise. Typically, 10-40 AFM measurements were made per each experimental condition. For AFM measurement of cells transduced with α-actinin, the microscope fluorescent mode was used to locate transduced cells with the GFP tag, followed by AFM measurements on those cells; for cell transduced with RhoA, the fluorescent tag was difficult to visualize in our AFM system, and thus this step was not taken before AFM measurements. However, transfection was separately confirmed using fluorescent microscopy and western blotting (22).

### OMTC Measurements

Cells were seeded to 96 well plates (Corning, Corning, NY) in the presence 10% serum. Ferromagnetic beads (4.5 μm in diameter), coated with poly-L-lysine (PLL), were allowed to attach to the cells for 20–30 min (14). The beads were then magnetized with a strong magnetic pulse in the horizontal direction and twisted with a much weaker oscillatory magnetic field (0.77 Hz) in the vertical direction. The torque on the bead was automatically adjusted to achieve a median bead translation of approximately 40 nm, and bead motion was quantified by image analysis. The ratio of magnetic torque to bead motion was used to determine complex shear modulus of cells and elastic modulus, *g’*, which has units of pascals per nanometer, was used as a measure of cell stiffness as previously described (14, 23).

### TM Measurements

Cells were seeded on collagen-coated acrylamide gels in 96 well plates or 35mm glass bottom dishes in the presence of 10% serum (media for each cell type as described above) throughout the experiments. The gels were prepared as previously described (24) and had a Young’s modulus of 8 kPa for SC cells and confluent MEFs and 2.4 kPa for sparsely seeded MEFs. A Leica epifluorescence microscope was used to take images of fluorescent microparticles on acrylamide gels. Based upon these images, displacements made by cells on gel were calculated using particle image velocimetry (25) and from the displacement fields, traction was retrieved using Fourier transform traction microscopy (25, 26). For confluent and superconfluent cells, root mean square (RMS) traction was used as a measure of average cellular contractile force (24) and for sparsely seeded cells, the contractile moment was used as a measure of average cellular contractile force (26).

### Confocal Imaging

Cells were seeded onto glass cover slips 48 hours prior to fixation and stained for confocal experiments. 100% methanol (Sigma, Milwaukee, WI) was used as the fixative for experiments where vimentin was the only cytoskeletal filament to be stained (Mendez et al. 2010). For all other experiments, 4% paraformaldehyde (Electron Microscopy Sciences, Hatfield, PA) was used (pH=7.4). Methanol fixation was done by incubating cells with precooled methanol for 10 minutes inside a −20°C freezer, and for paraformaldehyde fixation, cells were fixed for 10 minutes at room temperature. For cases where the secondary antibody was raised in goat (phosphorylated myosin and vimentin staining), samples were blocked in 10% normal goat serum (Life Technologies, Grand Island, NY) for 20 minutes to block for the non-specific binding prior to incubation with primary antibody. For all other cases, cells were incubated with Image-iT^®^ FX Signal Enhancer (Life Technologies, Grand Island, NY) for 20 minutes prior to incubation with primary antibody.

For dexamethasone experiments, cells were stained for F-actin (30 minutes incubation with Alexa Fluor^®^ 568 Phalloidin, Life Technologies, Grand Island, NY), vimentin (30 minutes incubation with Alexa Fluor^®^ 488 Conjugated Vimentin (D21H3) XP^®^ Rabbit mAb, Cell Signaling Technology, Danvers, MA), the nucleus (15 minutes incubation with Hoechst 33342 (1:10000), Thermo Fischer Scientific, Grand Island, MA), and phosphorylated myosin (overnight incubation at 4°C with Phospho-Myosin Light Chain 2, Ser19 (1:100) followed by two washes with PBS and one hour incubation with goat anti rabbit IgG (H+L) Alexa Fluor 488 (1:400), Cell Signaling Technology, Danvers, MA). For RhoA and α-actinin experiments, cells were stained for F-actin (30 minutes incubation with Alexa Fluor^®^ 568 Phalloidin), and the nucleus (15 minutes incubation with Hoechst 33342 (1:10000)).

A Zeiss 510 LSM inverted two-photon confocal microscope equipped with 10x (NA=0.3), 20x (NA=0.8), 40x (water immersion, NA=1.2), and 63x (oil immersion, NA=1.4) objective lenses and excitation sources of Ar (458, 488, 514 nm), HeNe (543 nm), HeNe (633 nm), 2-photon (690-1024 nm) was used to create confocal images from the samples (Carl Zeiss, Thornwood, NY). Thickness of MEFs and fluorescent intensity for the peripheral F-actin and P-myosin for SC cells were measured using Fiji image processing software.

Bead embedding depths were examined in wildtype and knock-out MEFs by using a confocal microscope (Leica TSC SP5, 63×/1.2-NA water immersion lens) after staining cytoplasm with a fluorescent dye (Cell-Tracker Green; Thermo Fisher, MA) (27). Using ImageJ, the percentage of bead embedding was quantified (28).

### Structured Illumination Microscopy (SIM)

MEFs were fixed with 3% paraformaldehyde, permeabilized with 0.1% Triton-X100 (Sigma Aldrich, Milwaukee, WI) in PBS for 10 minutes at RT, and then rinsed briefly in PBS. The samples were subsequently incubated in chicken anti vimentin antibody (1:2000, Biolegend, San Diego, CA) in PBS containing 5% normal goat serum (Jackson Immuno Research, West Grove, PA) for 30 minutes at RT followed by washing with PBS containing 0.05% Tween 20 twice for 3 minutes, and finally washed with PBS for 3 minutes. The slides were then incubated in goat anti chicken antibody (Alexa Fluor^®^ 488, 1:400, Life Technologies, Grand Island, NY) and Alexa Fluor^®^ 568 Phalloidin (1:400, Life Technologies, Grand Island, NY) for 30 minutes. Slides were then washed with PBS containing 0.05%Tween 20 twice for 3 minutes, and finally washed with PBS for 3 minutes. The coverslips were mounted using Prolong Diamond (Thermo Fischer Scientific, Grand Island, MA). 3D-SIM was carried out with a Nikon Structured Illumination Super Resolution Microscope System (Nikon N-SIM; Nikon, Tokyo, Japan) using an oil immersion objective lens CFI SR (Apochromat TIRF100× 1.49 NA; Nikon).

### Statistical Analysis

Because measurements of cell stiffness are not normally distributed (11, 12, 29, 30), data were logarithmically transformed for statistical analysis and determined as geometric means ± standard error around the geometric mean (15) for each cell strain, unless otherwise indicated; averages taken over cell strains were arithmetic. Linear regression was used to examine the relationship between drug concentration (dexamethasone) and log cell stiffness or log cell traction. ANOVA was used to compare the effects of inductions of GFP, α-actinin or RhoA on log cell stiffness and to compare vimentin KO MEFs with controls, allowing for unequal variance between groups, and using Tukey to account for multiple comparisons. JMP was used for these analyses with a significance level of 0.05; the minimum reported p-value is 10^−4^. Details of which JMP routines were used are found in the Supplemental Information.

### Finite Element Modeling (FEM)

FEM was used to model the influence of the cell cortex on AFM and OMTC measurements of cell modulus following the approaches of Vargas-Pinto et al. (12) and Mijailovich et al. (31), respectively. The cell was modeled as a cylinder of radius 10 μm (AFM) or 22.5 μm (OMTC) and a range of cell thickness from 2.75-15 μm with a nominal thickness of 5 μm. The radius of the cell was chosen to be sufficiently large such that the strains at the edge of the domain were less than 0.1% of the maximum strain near the probes, and a 50% increase in radius did not significantly affect the results (<0.1% difference in all parameters for AFM and <0.13% for OMTC). Except for very small or narrow cells, the assumed shape of the cells would therefore have little effect on the results.

The cortex of the cell was on its upper surface and had a thickness of 400 nm (12) in all cases with no-slip between the cortex and internal cytoskeleton (31); we have previously examined the effect of variations in cortex thickness and found that the modeling results are not particularly sensitive to this parameter (12). The intracellular cytoskeleton and cortex are each modeled as homogeneous elastic materials with the cortex having a modulus that ranged from 1 to 100 times the modulus of the intracellular cytoskeleton. Although the material was assumed to be incompressible, a Poisson’s ratio of 0.49 was used for numerical stability. The AFM round-tips and OMTC probes were modeled as rigid spheres (12, 31). The bottom surface of the cell for both AFM and OMTC was pinned for zero displacement; the other surfaces were stress-free except where the probes contacted the cell as discussed below. Stress, strain and deformation fields were determined using ABAQUS/CAE 6.13 finite element software (Simulia, Providence, RI) with an explicit solver (32). An apparent modulus (*E_apparent_*) of an equivalent single layer, homogeneous cell was computed that would deform to the same extent as the two-layer cell model for the same external torque (OMTC) or force (AFM)

#### Model of AFM Tip

A 2-D axisymmetric model was used (12). The indentation is assumed to be frictionless, consistent with the Hertz model (12), and the apical surface of the cortex was assumed to remain in contact with the AFM tip throughout the deformation process (33). The diameter of AFM rounded probes investigated varied from 0.8 μm to 10 μm and had an indentation into the cell of 400 nm. For the AFM sharp-tip, a conical shape was assumed with a tip diameter of 40 nm and a tip half angle of θ=20°, measured from a scanning electron micrograph of a typical sharp tip, or 36° (nominal value reported by the manufacturer). Indentation of the sharp-tip into the cell was limited to 80 nm as higher indentations caused excessive numerical element distortion and even with remeshing, the model did not converge to a solution, a limitation noted in other such studies (12, 34). This is also the reason that round-tip probes smaller than 0.8 μm in diameter could not be modeled for a 400 nm indentation. We note that the behavior for progressively smaller round tips, indenting 400 nm into a cell, are consistent with our results for a sharp tip indented 80 nm into a cell (e.g. Figures 6, S4, S5). To calculate E_*apparent*_ for AFM, the reaction force for the given indentation of the AFM tip into the cell were determined using ABAQUS for each case and matched to that of a homogenous case (cortex and internal cytoskeleton with equal modulus) that gave the same force for the same indentation.

#### OMTC Model

For OMTC modeling, a full 3D geometry model was used. A no-slip boundary condition was applied between the probe (4.5 μm in diameter) and cell cortex(31). The magnetic field applied in OMTC was modeled as a torque per unit volume (60 Pa (31)) applied to the center of the bead causing the bead rotate around its central axis. Depth of the bead embedding was varied from 10-50% of the bead diameter (35, 36). To calculate E_*apparent*_ for each case, the horizontal displacement of the OMTC probe for the given torque was determined using ABAQUS and matched to that of a homogenous case (cortex and cytoskeleton with same modulus) that had the same torque. To validate our model, we followed Mijailovich et al. (31, 32) and defined a parameter β that allowed an effective shear modulus (G_d_=βG) to be determined based on bead embedding depth:

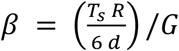

where *T_s_* is the torque applied per unit bead volume, *R* is the bead radius, *d* is the bead linear translation caused by the applied torque and *G* is the shear modulus of the homogenous medium in which the bead is embedded. *β* can be thought of as the effective stiffness of the cell (torque required for a given bead rotation) at a given bead embedding depth relative to the stiffness that would be seen for a homogenous cell with *E_cortex_*=*E_intracellular_* and a bead embedding depth of 50% in an infinite medium. We found values of *β* that were in good agreement with those found by Mijailovich et al. for the case they examined (*E_cortex_*=*E_intracellular_*) (see Fig. S10).

## RESULTS

### Alterations induced by Dexamethasone, α-actinin and Rho-A

The monolayer of endothelial cells lining the inner wall of SC in the eye is subject to a pressure gradient, and the mechanical properties of these cells have been shown to be important to their biological barrier function (15). Using AFM, TM and OMTC, here we further characterized the mechanical properties of these cells before and after treatment with dexamethasone, or transduction with adenovirus encoding for Rho-A or α-actinin. Dexamethasone can cause glaucoma and is known to generate cytoskeletal changes in SC cells (37); Rho-A is a cytoskeletal regulator (38) and α-actinin is an actin cross-linker (39).

Figs. 1 and S11 shows the distribution of filamentous actin (F-actin) and phosphorylated myosin light chain (P-myosin) in confluent SC cells treated with dexamethasone (1μM) as compared with controls. In both control and treated cells, we found stress fibers distributed throughout the cells, particularly along the cell peripheries (Fig 1A, D). The distribution of P-myosin was similar to that of F-actin (Fig 1B, E). Compared to control treatment, we found that dexamethasone-treated cells had denser stress fibers and P-myosin at the periphery of the cells. Fluorescent intensity measurements confirmed these conclusions showing that the dexamethasone-treated group (1 μM) had higher cortical F-actin (p<10^−4^) and P-myosin (p=0.0002) as compared with control.

**Fig. 1:**
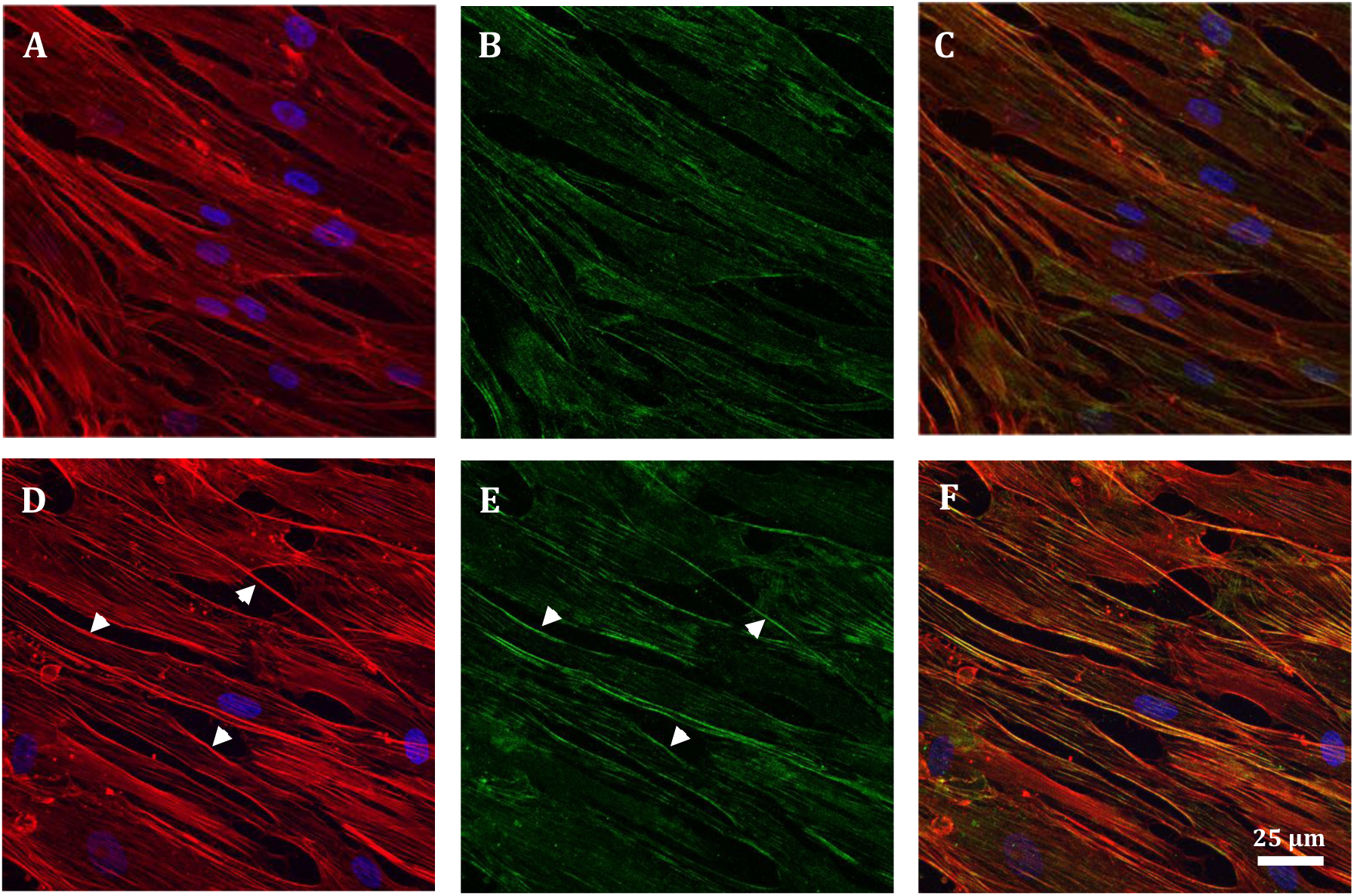
Confocal images of confluent monolayer of vehicle only control (top row) as compared with 1μM dexamethasone-treated (bottom row) SC cells; F-actin(red), phosphorylated myosin (green), nucleus (blue). Cortex and stress fibers are seen in both groups but the cortex is more prominent in treated SC cells (white arrows in D). Also, while p-myosin is present in both groups, it is more concentrated at the cortex of treated cells (white arrows in E). (C) and (F) are overlaid images.

Increasing concentrations of dexamethasone caused dosage-dependent increases in both SC cell stiffness (with sharp-tip AFM) and cell traction (TM), respectively (Fig. 2A; linear regression: p<10^−4^). In contrast, dexamethasone had either no effect on SC cell stiffness as measured with AFM round-tips (p≈1) but a small statistically significant decrease in cell stiffness as measured with OMTC. (Fig. 2B; p<10^−4^)

**Fig. 2:**
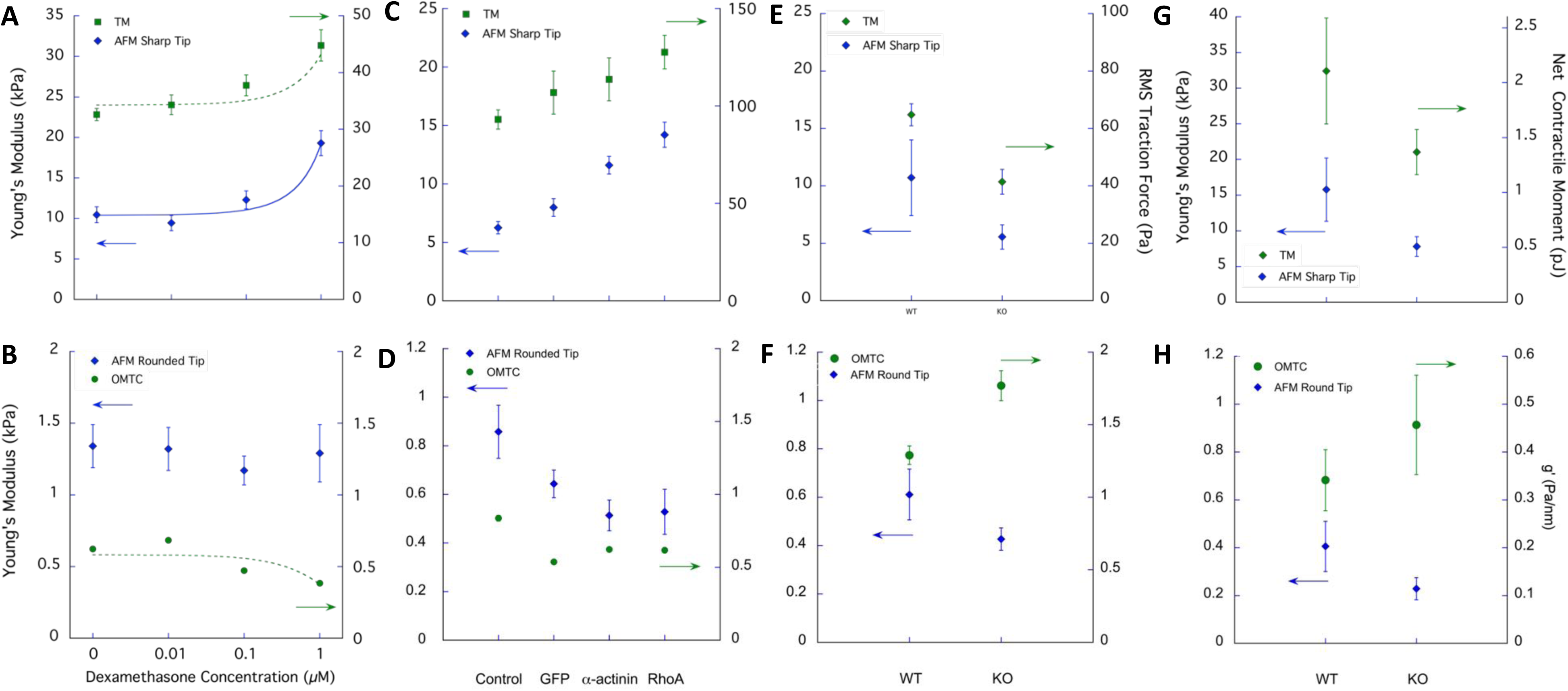
Measurements of cell biophysical properties of SC cells after (A, B) treatment with varying concentrations of dexamethasone as compared with vehicle alone, or (C, D) overexpression of α-actinin or RhoA as compared with GFP or control. (E-H) shows comparison of wildtype (WT) MEFs with vimentin knockouts (KO). Super confluent monolayers were used for the dexamethasone studies on SC cells (A,B), confluent for the α-actinin or RhoA expression studies on SC cells (C-D), and both confluent (E-F) and sparsely seeded cells for the MEF studies (G, H), as described in the Methods section. Geometric mean ± S.E. about geometric means. Curves shown in (A, B) are best regression fit to the data; the equations are given in Supplementary Information. Data for individual cell strains are founds in Supplementary Figs. S2 and S3.

Fig. 3 shows the distribution of F-actin in SC cells before and after transduction with green fluorescent protein (GFP: a control), or overexpression with α-actinin, or RhoA. Cells in the GFP group (3B) had an F-actin distribution similar to that of control cells (3A). However, cells transduced with either α-actinin and RhoA showed an altered morphology with thicker and more prominent cortical stress fibers (arrows in Figs 3C, D).

**Fig. 3:**
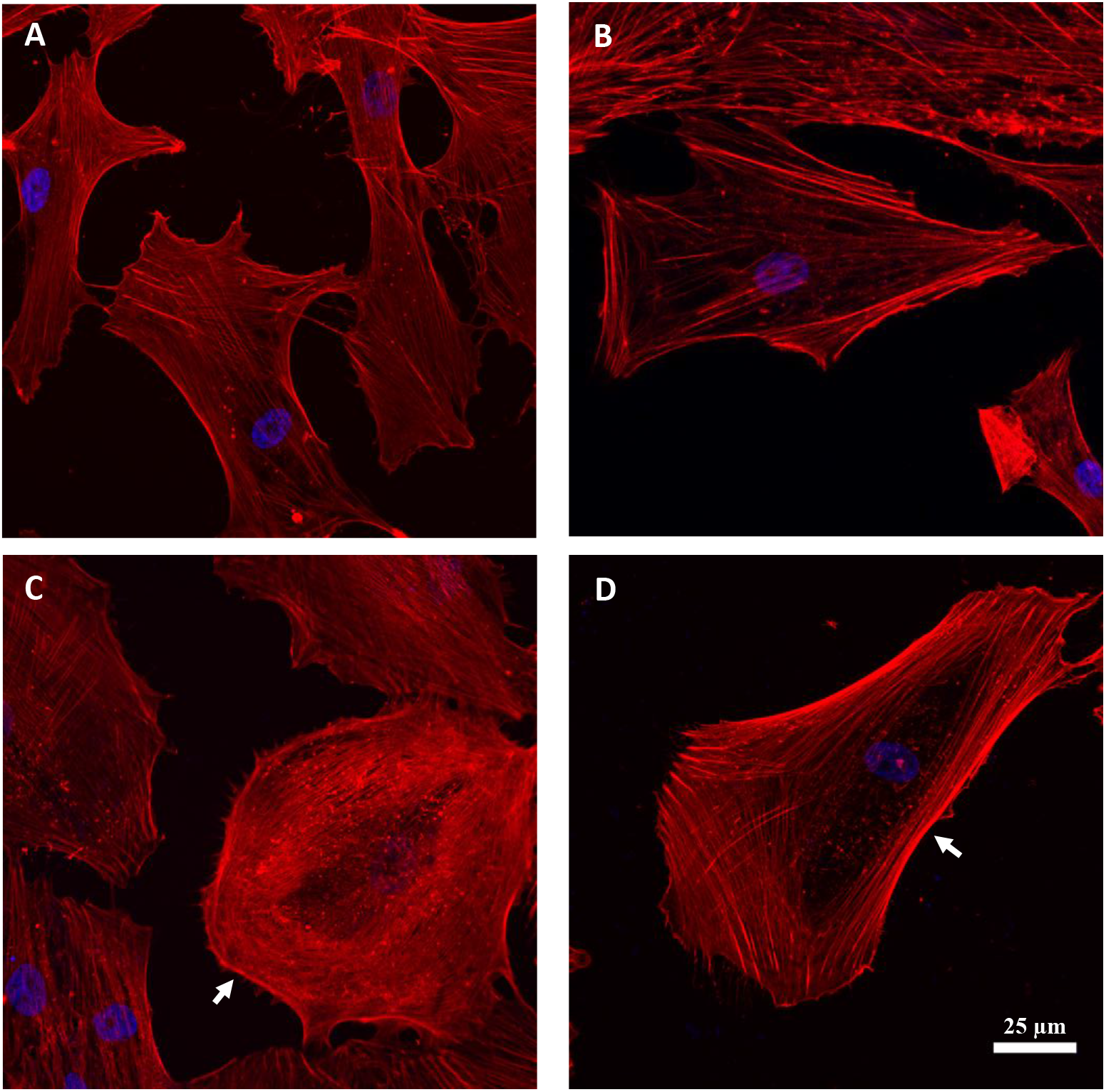
Confocal images of SC cells. (A) WT-non-transduced, (B) GFP-transduced, (C) α-actinin-transduced, and (D) RhoA-transduced; F-actin (red), nucleus (blue). Control and GFP groups had similar F-actin distribution and cortex structure (A, B). The α-actinin-transduced cells showed an altered morphology and had a relatively thicker cortex (white arrow in C). The RhoA-transduced cells showed significant accumulation of stress fibers at peripheral regions and a more prominent cortex (white arrow in D).

Comparison of cell stiffness in wild type (WT) and GFP-transduced cells using AFM with a sharp-tip showed no difference (p>0.8). However, overexpression of both α-actinin (p<2×10^−4^) and RhoA (p<10^−4^) increased cell stiffness compared to control cells or cells transduced with GFP (Fig. 2C). Using TM, we found traction to be increased after transduction of RhoA (p<10^−4^) compared to GFP transduced cells and controls (Fig 2C). Due to one non-responsive cell strain (Fig. S3C), the increased traction after transduction of α-actinin was not statistically significant (p>0.3). In contrast, measurements with both AFM round-tips and OMTC showed that overexpression of α-actinin and RhoA decreased cell stiffness compared with control cells (AFM: p<0.03; OMTC: p<10^−4^) (Fig. 2D). When compared with the GFP group, AFM round tips revealed a trend for decreased cell stiffness for the α-actinin and RhoA overexpression groups while OMTC revealed a trend for increased cell stiffness, but in neither case was the difference significant (Fig 2D). OMTC also showed that GFP decreased cell stiffness relative to control cells (p<0.001); this trend was also apparent with AFM round-tips but was not statistically significant (p>0.4).

### Vimentin knockout reduces cell stiffness and traction

Vimentin is a Type III intermediate filament protein that assembles into a major cytoskeletal system (40–42). To study the role of vimentin in cytoskeletal mechanics, we used immortalized mouse embryonic fibroblasts (MEFs) isolated from a mouse knockout of the intermediate filament gene encoding vimentin (19). In vimentin KO MEFs, there was no vimentin, as expected, and in WT MEFs there was a robust network of vimentin intermediate filaments throughout cell body (Fig. 4C; 4D). In WT cells, vimentin filaments were mostly found in the intracellular region but also within close proximity to the actin-rich cortex (Fig. 4E-G). No obvious difference was seen in the F-actin distribution between the WT and vimentin-KO MEFs (Fig. 4A, B).

**Fig. 4:**
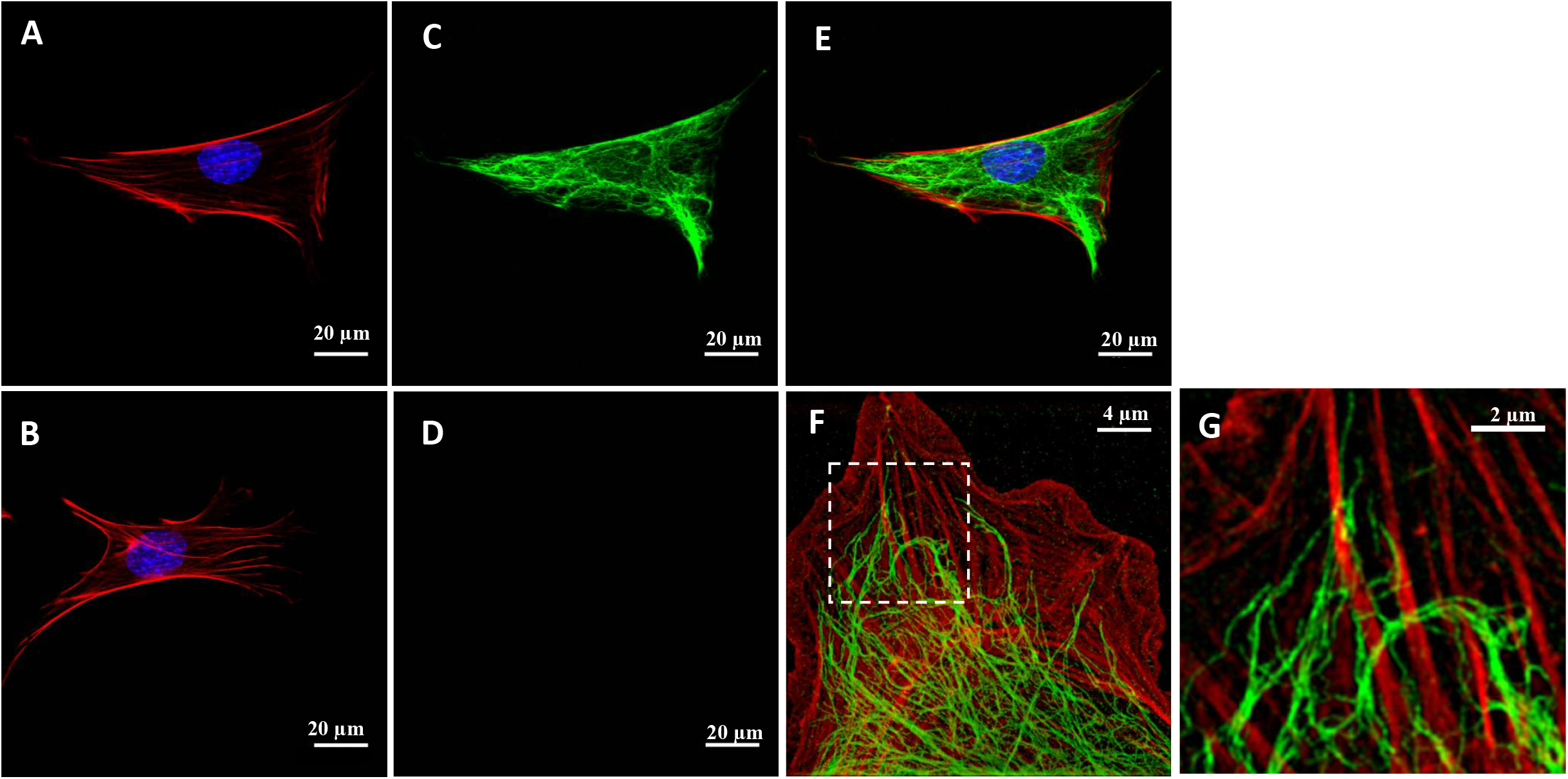
Confocal images showing F-actin (red), nucleus (blue) (A, B) and vimentin intermediate filaments (green) (C, D) of WT MEFs (A, C) and vimentin-KO MEFs (B, D). Panel D shows no staining in a vimentin-KO MEF. Panel (E) shows merged image of A and B. Panel F is structured illumination micrograph (SIM) of a wildtype MEF at the basal cortex level showing the close association between actin stress fibers and vimentin intermediate filaments. Magnified image of the insert in (F) showing vimentin intermediate filaments surrounding and interconnecting actin stress fibers (G).

We compared the mechanical behavior of vimentin-KO MEFs to WT MEFs using both confluent and sparsely confluent seeding conditions. Vimentin-KO MEFs were generally softer than WT MEFs (using AFM sharp tips; confluent monolayers, p<0.03, Fig 2E; sparsely cultured p<0.06: Fig 2G) and exerted less traction (using TM; confluent monolayers, p<0.03, Fig 2E; sparsely cultured p<0.03, Fig 2G). Using AFM round-tips, we also found that vimentin-KO MEFs were softer than WT MEFs, but the differences did not reach statistical significance for either confluent monolayers (p<0.3) or sparsely cultured cells (p<0.07) (Fig 2F, 2H).

Unexpectedly, when we measured stiffness using OMTC, the vimentin-KO MEFs were stiffer than WT MEFs (confluent monolayers p<0.0001; sparsely cultured cells p<0.02) (Fig 2F, 2H). These observations were in contrast to the measurements taken using AFM sharp-tips, AFM round-tips and TM. To understand why the results obtained by OMTC differed from those obtained by AFM and TM in MEFs, we measured the bead embedding depth and cell thicknesses, both of which impact stiffness measured by OMTC (31). We found that the bead embedding depth was similar in both cell types with no significant difference between WT and vimentin-KO MEFs (mean of 63%, n=14-18; p>0.6). However, the thickness of the vimentin-KO MEFs (3.2±0.4 μm, mean±SD; n=20) was significantly smaller than that of the WT MEFs (4.2±0.5 μm, mean±SD; n=30) (Fig 5) (p<10^−4^). As shown below, finite element modeling reconciles these observations.

**Fig. 5:**
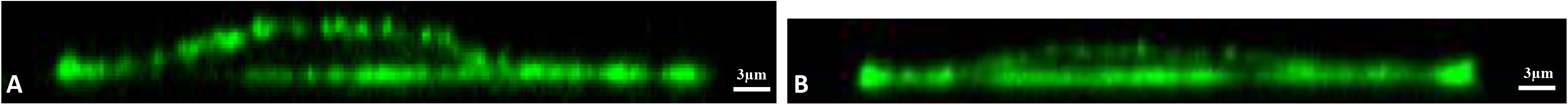
Confocal cross-sectional representative images of typical WT (A) and vimentin-KO MEFs (B) show that the vimentin-KO MEFs are thinner than the WT MEFs.

### Finite element modeling (FEM) shows that the cell cortex differentially affects differing measurements of cell stiffness

To model OMTC, we followed the approach of Mijailovich et al. (31) but allowed for a cell cortex that was significantly stiffer than the underlying intracellular network. To model AFM, we extended the approach of Vargas-Pinto et al. (12) to determine the effect of AFM tip size. We modeled a cell as a thin cylinder with a nominal thickness of 5μm that had two layers: a 400 nm thick cortex and an underlying intracellular network. Embedded in the top of the cell was either a sharp or round AFM tip that exerted a vertical force on the cell, or a round OMTC probe that exerted a torque on the cell. Details are in the Methods section.

We first examined the strain energy density distribution caused by AMF tips or the OMTC probe when the cortex has the same stiffness (*E_cortex_*) as the intracellular cytoskeleton (*E_intracellular_*) and when it is 50 times stiffer (Fig. 6; Supplemental Information). For the AFM sharp-tip, strain energy is largely confined to the cortex (Figs 6A,B), consistent with the previous findings (12). For the smallest AFM round-tip probe (0.8 μm), strain energy is also largely confined to the cortex but also extends appreciably into the intracellular network, particular for the case of a stiff cortex (Figs. 6C,D). For the largest AFM round-tip probe (10 μm), strain energy extends throughout the cell beneath the probe, but is somewhat broader for the case of a stiff cortex (Figs. 6E,F). Finally, for OMTC the strain energy distribution is intermediate between the cases of the small and larger AFM tips, but is more similar to the latter, with the strain energy spread out more broadly for the case of a stiff cortex (Figs. 6G,H). These results suggest that when the size of probe exceeds the thickness of cortex, strain energy spreads out of the cortex, particularly when the cortex is much stiffer than the intracellular network.

**Fig. 6:**
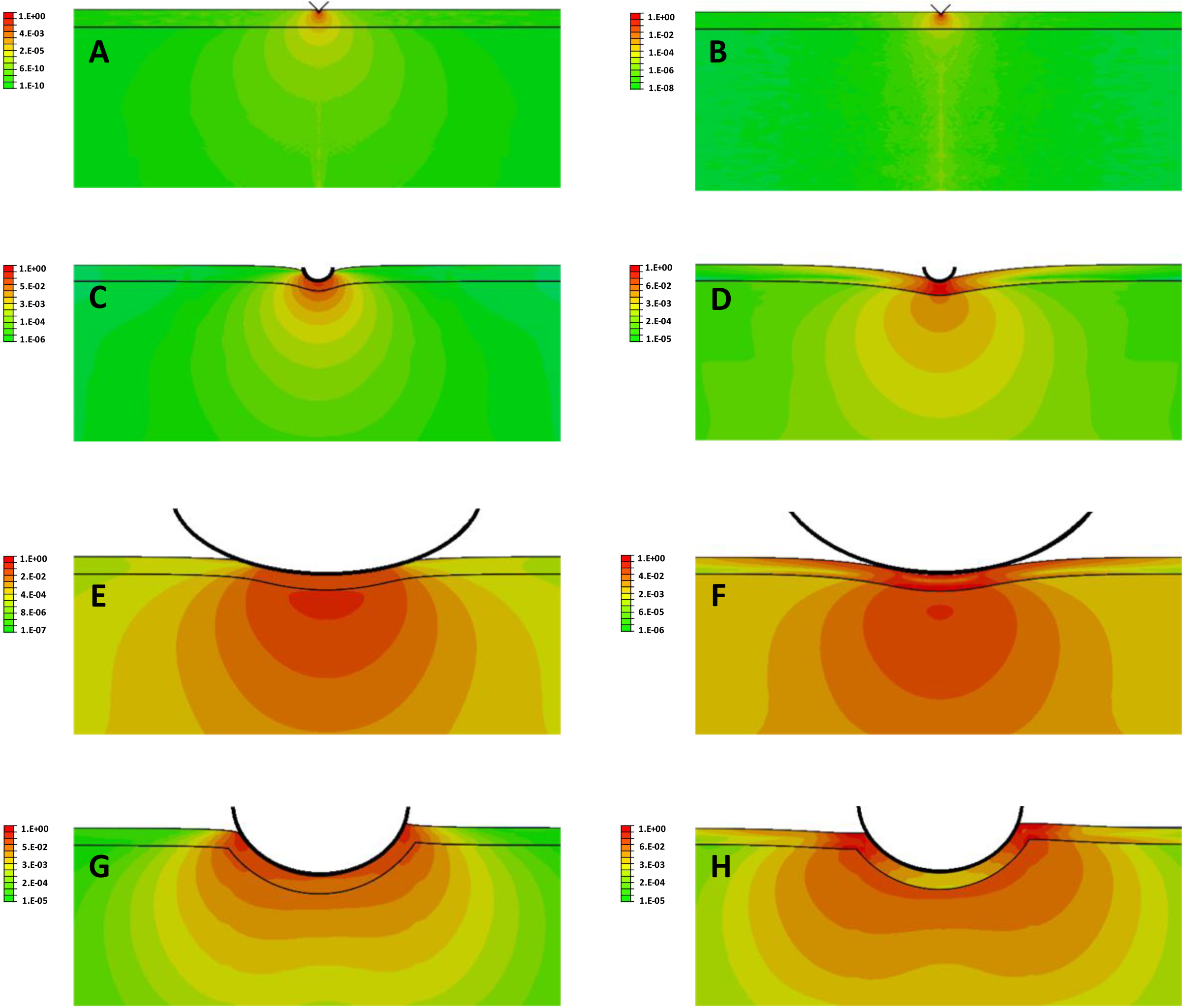
Strain energy distribution (log scale) for indentation into a cell of a sharp AFM tip (36° half angle) (A,B), a 0.8 μm diameter rounded AFM tip (C, D), a 10 μm diameter rounded AFM tip (E,F) and a 4.5 μm OMTC bead (G,H). The indentation is smaller in cases A, B (80 nm), due to numerical limitations as discussed in the Methods section, than for the other AFM tips (400 nm). The OMTC bead is embedded 25% of its diameter into the cell and twisted by a torque of 60 Pa applied in a counter-clockwise fashion. Panels A, C, E, G are for cases with *E_cortex_*=*E_intracellular_*; panels B, D, F, H are for *E_cortex_*=50\**E_intracellular_*. The cortex in each panel is the narrow region between the two horizontal black lines and has a thickness of 400 nm prior to indentation. The strain energy distribution in each panel is normalized to the maximum strain energy in that panel and a log scale is used. Cell thickness is 5 μm.

The effect of cortex stiffness on measurements by AFM and OMTC was quantified by computing an apparent modulus (*E_apparent_*) for an equivalent single layer, homogeneous cell that deformed to the same extent as the cell model with cortex, as a function of *E_cortex_*/*E_intracelluar_* (see Methods). For AFM sharp-tips, *E_apparent_* ranged from 30~50% of *E_cortex_* (Fig 7A inset) while for AFM round-tips, *E_apparent_* was less than 10% of *E_cortex_*. For an AFM tip diameter of 10 μm, *E_apparent_* was nearly independent of *E_cortex_* and near the value of *E_intracelluar_* (Fig 7A).

**Fig. 7:**
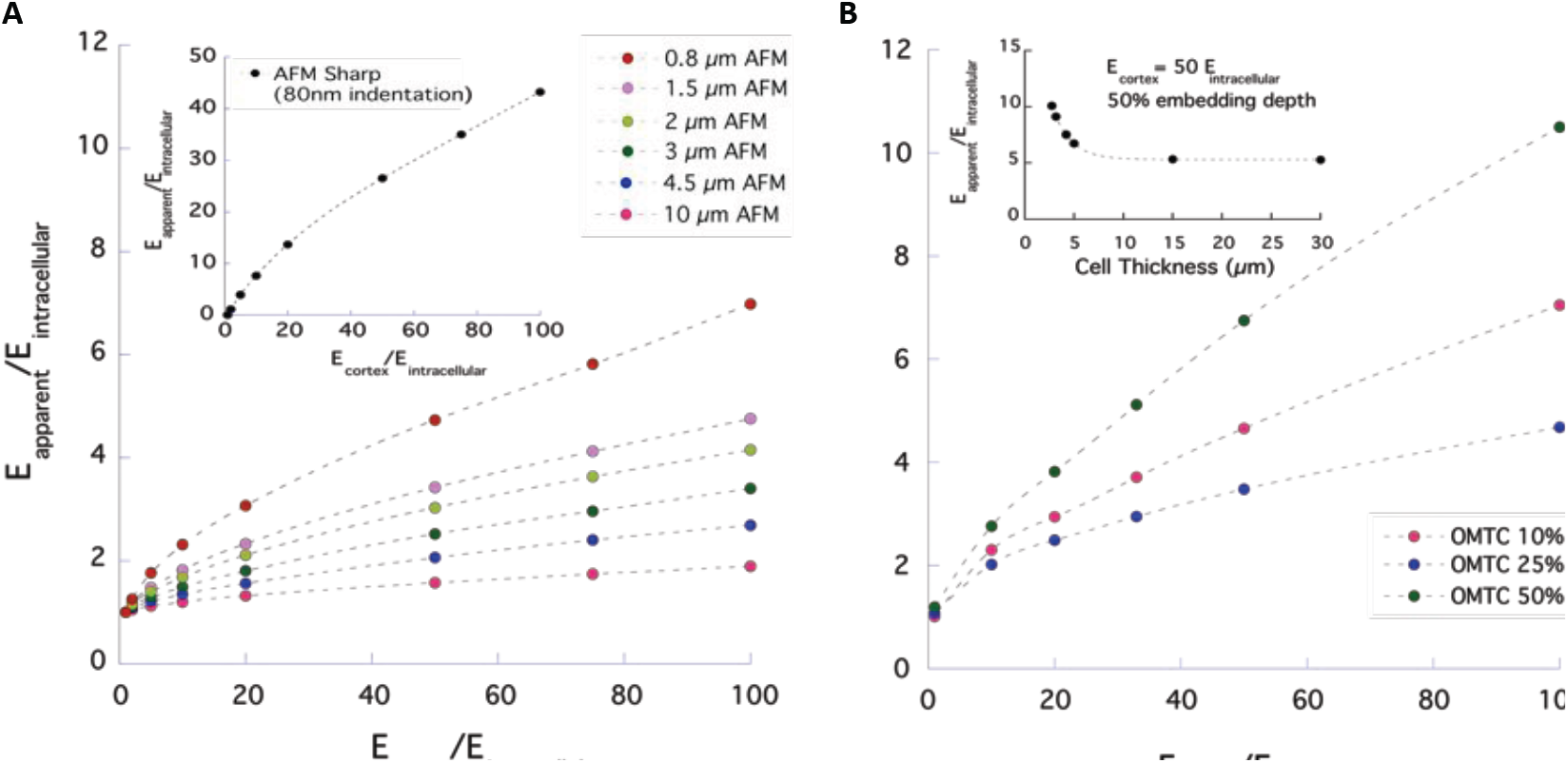
(A) *E_apparent_*/*E_intracellular_* as a function of *E_cortex_*/*E_intracellular_* for an AFM round-tip of diameter 0.8 μm to 10 μm with an indentation of 400 nm. Inset shows result for AFM sharp-tip with indentation of 80 nm; θ/2=20°. Cell thickness is 5 μm. (B) *E_apparent_*/*E_intracellular_* as a function of *E_cortex_*/*E_intracellular_* for an OMTC probe of diameter 4.5 μm for embedding depths of 10, 25 or 50% of bead diameter. Cell thickness is 5 μm. Inset shows results for an OMTC probe embedded 50% into a cell with *E_cortex_*/*E_intracellular_*=50 for cell thicknesses ranging from 2.75-30 μm.

For OMTC the predictions were similar to those of AFM round-tips, with *E_apparent_* much closer to *E_intracelluar_* than to *E_cortex_* (Fig. 7B). These results were very different than those of AFM sharp-tips and show the stiffness measured by OMTC is largely characteristic of the intracellular network although it is still somewhat affected by cortex stiffness. Interestingly, embedding depth does not have a monotonic effect on *E_apparent_* but decreases with embedding depth for shallow embedding and then reversing this trend for deeper embedding (Figs. 7B, S7). This is caused by the distal cortex (the cortex that is not surrounding the bead) that becomes somewhat more important for deeply embedded beads (Fig. S8), and by a substrate effect (insert in Fig. 7B; Supplemental Information).

### The effect of decreased KO MEF cell thickness on OMTC and AFM measurements

Our OMTC measurements suggested that vimentin-KO MEFs were stiffer than WT MEFs, in contrast with the measurements using the other techniques (Fig 2E-H). Because we found that vimentin-KO MEFs are thinner than WT MEF, we examined whether this thickness difference might explain the discrepancy, particularly in the context of deeply embedded OMTC beads. For cells thinner than 5 μm, the modulus rises rapidly with decreasing cell thickness (inset Fig. 7B; Fig. S6), which is a behavior that would explain why the OMTC showed KO MEFs to be stiffer than WT. The effect of decreased thickness of KO MEFs on *E_apparent_* (Fig. S9) would also explain why the AFM round-tip results were lower in KOs than WTs but did not reach statistical significance.

## DISCUSSION

Here we have examined how the choice of surface probe impacts the measurement and interpretation of cell biomechanical properties. In response to a panel of interventions, and in both cell types investigated, cell stiffness measured using AFM sharp-tips and contractile forces measured using TM tracked similarly to one another (Fig. 2A, C). However, these measures did not track with cell stiffness measured by OMTC or with AFM round tips. To explain these differential responses, we turned to FEM simulations. Simulations of AFM round tips (5 μm or larger) and OMTC were sensitive to stiffness of the intracellular network but relatively insensitive to stiffness of the cell cortex (Fig. 7A,B); these results also showed that OMTC is particularly sensitive to cell thickness. Simulations of AFM sharp tips, by contrast, were more sensitive to the stiffness of the cell cortex with values of E_*apparent*_ comparable to cortex stiffness (Fig. 7A inset). Taken together, these findings suggest that measurements using AFM sharp tips and TM tend to emphasize biomechanical properties of the cell cortex, whereas measurements using OMTC and AFM round-tips tend to emphasize stiffness of the intracellular network.

OMTC measurements on cells previously have been suggested to characterize cortical stiffness (11, 18, 27, 43). The relative insensitivities of OMTC to cortex stiffness found in our simulations was due to the large size of these probes as compared to the thickness of the cortex leading to substantial strain energy in the intracellular network. Our finding that OMTC is sensitive to intracellular network stiffness is consistent with studies of molecular crowding caused by cellular volume reduction showing increased cell stiffness as measured using OMTC but no change in cellular contractile force as measured by TM (23, 27).

We note that the FEM used here greatly simplifies cytoskeletal mechanics by treating the cortical and intracellular networks as two homogeneous linearly elastic materials. The goal of these models was not to replicate the complex geometry of the cytoskeleton but instead to evaluate how a stiff cortex may affect measurements of cell stiffness using different probes. Since these measurements are conventionally interpreted using models of cells that are homogeneous and linearly elastic (21, 31, 44, 45), examining the effect of a stiff cortex on these stiffness measurements necessarily required use of a similar model, modified to include a cortex. A more realistic FEM model of the cell would likely further strengthen our conclusions, since the intracellular cytoskeletal network is connected to the cortical network, and this will induce propagation of apically applied force/torque from OMTC beads or larger AFM rounded probes to deeper cytoskeletal structures or even basal regions of the cell (46, 47).

Vimentin is a structural protein that polymerizes into a major intermediate filament cytoskeletal system (40, 48–50). Our imaging studies showed a network of vimentin intermediate filaments in WT MEFs that were entirely absent in vimentin-KO MEFs (Fig. 4C,D). We also saw evidence of vimentin in the actin-rich cortical region of the WT MEFs (Fig. 4 F,G), in agreement with previous findings in chicken embryonic fibroblasts (51) and bovine pulmonary artery endothelial cells (52). We used AFM sharp tip and TM to measure cell stiffness and contractile state of the cortex of the cells and found both significantly reduced in vimentin-KO MEFs as compared to WT MEFs (Fig. 2E,G) leading us to conclude that vimentin intermediate filaments contribute to or influence cortical stiffness and cellular contractile force. Measurements using AFM round tips allowed us to conclude that the intracellular network of vimentin-KO MEFs also has reduced stiffness as compared to WT MEFs (Fig. 2F,H).

Guo et al. (18) used optical tweezers and found that the cytoplasmic stiffness of vimentin-KO MEFs was reduced as compared to WT MEFs, consistent with our finding of reduced intracellular stiffness. They also used OMTC and concluded that vimentin does not influence cortical stiffness, in contrast to our AFM sharp tip and TM findings. However, we found decreased thickness in vimentin-KO MEFs relative to WTs (Fig. 5), consistent with a previous report (53), and were able to show that these cell thickness differences would substantially impact OMTC measurements of cell stiffness (Fig. 7B, insert), leading us to conclude that OMTC measurements of vimentin-KO MEFs stiffness need to be re-considered in light of the results reported here.

Wang and Stamenovic (54) found that at low levels of strain, the stiffness of WT and vimentin-KO MEFs as measured using OMTC were similar to one another, while at higher strain levels, the stiffness of WT cells was greater than that of vimentin-KO cells. Mendez et al. reported similar results using AFM round-tips (55), and these investigators concluded that the vimentin network contributes to cell stiffness when the cytoskeleton is under high levels of strain. Our studies are consistent with this finding, showing that WT cells had a higher cell stiffness than vimentin-KOs when measured with sharp AFM tips that generate high levels of strain, as compared with AFM round-tips and OMTC that generate much lower levels of strain (Figs. 2, S4).

Our studies focused on examining regional differences in cytoskeletal structure and how these differences affect measurements of cell mechanics by external probes. We examined the effect of several agents that were anticipated to affect cellular mechanics and found that they affected the cortical regions differently than the intracellular network. RhoA regulates cytokinesis (3) causing stiffening of the cell cortex (3) and retraction of the vimentin intermediate filament network (56–58). This is consistent with our findings that RhoA overexpression increases cortical cell stiffness and traction as measured by AFM sharp-tips and TM, respectively, and somewhat reduces intracellular network cell stiffness as measured by AFM round-tips (see Fig. 2). This may also relate to our finding that α-actinin overexpression causes similar changes (59, 60). These results also show that over-expression of regulatory factors such as RhoA and α-actinin can be transduced into cells to alter their cortical stiffness.

We also found that dexamethasone caused an increase in the cortical stiffness of SC cells (Fig. 2A), along with increased density of cortical F-actin and P-myosin (Fig. 1) with little effect on the intracellular network (Fig. 2B). Dexamethasone induces RhoA activation (61) and is associated with elevated stress fiber formation in human trabecular meshwork cells (62, 63), decreased hydraulic conductivity of SC and trabecular meshwork cell monolayers (64), and increased traction in alveolar epithelial cells (65). This may be related to the vasoconstriction that steroids are known to promote (66). Corticosteroids such as dexamethasone are widely used to offset allergies, asthma, and skin rashes, but their ocular use is known to elevate intraocular pressure in some individuals and can cause glaucoma (67). Our studies here, when considered with our previous work showing glaucomatous SC Cells to have increased stiffness (15), provide a mechanism by which dexamethasone could cause the elevated intraocular pressure characteristic of glaucoma.

Our AFM measurements, combined with FEM, suggest that the stiffness of the intracellular network is roughly 0.25-1 kPa (Figs. 2B, 2D, 2F, 2H, 7A) while that of the cortical cytoskeleton is at least one order of magnitude higher (Figs. 2A, 2C, 2E, 2G, 7A insert). As expected, both values are much higher than the reported stiffness of cytoplasm (0.005-0.01 kPa) measured using optical tweezers (18), and thus no conclusions can be drawn from our studies with respect to the cytoplasm (11). Table 1 shows a summary of our findings regarding the behavior of the different measurement techniques on the cells considered in this study. A recent report by Wu et al (68) using MCF-7 cells is consistent with these findings.

**Table 1.** Summary of Findings from Experimental Measurements and FEM

| **Technique** | **Typical Modulus<sup>+</sup>/Stress<sup>\*</sup>** | **Measurement Locus** | **Sensitivity to cell thickness** |
| --- | --- | --- | --- |
| AFM Sharp Tip | 5-25 kPa<sup>+</sup> | Cortical | insensitive |
| AFM 1.5 μm Rounded Tip | 0.5-4 kPa<sup>+</sup> | Cortical and intracellular | relatively insensitive |
| AFM 10 μm Rounded Tip | 0.25-1.5 kPa<sup>+</sup> | Intracellular network | sensitive |
| OMTC | 0.5-2 kPa<sup>+</sup> | Cortical and intracellular | sensitive |
| TM | 0.02-0.5 kPa<sup>*</sup> | Cortical | not investigated |

Our studies show that measurements of cell mechanics by different external probes are differentially sensitive to regional differences in cytoskeletal structure. While stiffnesses measured by AFM sharp- tips characterize the stiff cortical region of the cells, which is under active traction, similar trends are observed in cellular contractile force measurements by TM; this suggests that cells modulate both cortical stiffness and traction force through a similar mechanism. In contrast, AFM round-tips and OMTC are both less influenced by the cortex and instead probe the stiffness of the non-cortical intracellular network. These interpretations are supported by our FEM studies, which also demonstrate that probe sensitivity to cell thickness can explain the OMTC trends. By considering the regional emphasis of each probe type, our results additionally show that vimentin intermediate filaments play significant mechanical roles in both the intracellular and cortical domains of cells. Taken together, these results highlight the importance of probe choice in the interpretation of cellular mechanics measurements.

## AUTHOR CONTRIBUTIONS

A.V., M.J., C.Y.P, K.P., W.D.S., and C.Y.P. designed the experiments. A.V., C.Y.P., Z.Z., E.K.D. and H.W. performed the experiments. A.V., M.J., C.Y.P., J.J.F. and D.W. designed and evaluated the model; A.V. implement the model and performed the simulations. K.P., W.D.S. and R.G. supplied the cell lines. A.V., M.J., C.Y.P, J.J.F., R.G., K.P., W.D.S., D.W. and H.W. analyzed the data and wrote the manuscript.

## ACKNOWLEDGEMENTS

We acknowledge support from NIH EY019696 (MJ, WDS, MJ, JJF), P01HL120839 (JJF and DAW), P01GM096971 (RDG, DAW) and U54 CA193419 (RDG), Northwestern NUANCE Center, Center for Advanced Molecular Imaging, and the National Natural Science Foundation of China (11472062, ZZ).

## Supplemental Information

### AFM measurements

To determine the aggregate cell modulus (E) for a rounded AFM tips of radius R, we used the following relationship (69):

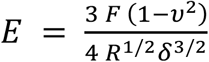

where F is the applied force, *δ* is the indentation, and v=0.5 is Poisson ratio (33, 70, 71). For sharp tips, we used a model for pyramidal indenters that accounts for the spherical cap at the apex of the AFM tip (21, 72):

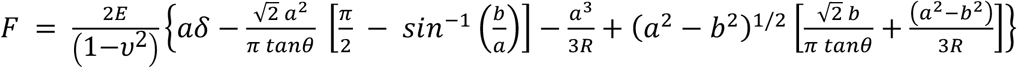

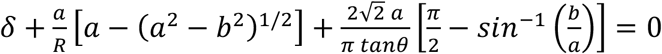

where *b*=*R* cos*θ* and *R* is the radius of the spherical cap (20 nm). Equation (S2b) is used to find the effective radius of contact *a* for any indentation *δ*, and then used in equation (S2a) to find *E*, for a given force *F*. We have previously shown that this model gives value of *E* that are relatively independent of *δ* for value of *δ* greater than approximately 100 nm (12). For the AFM sharp tips, we used a tip half angle of θ=20.

Fig. S1 shows typical force-ramp AFM curves and their corresponding fits from equations (S2) or (S1) for sharp (A, C) and rounded (B, D) AFM tips, respectively. The fits are found by determining the average modulus that best fits the data for *δ* ≳ 100 nm and then using that modulus in equation (S1) or (S2). The particular examples shown compare results for (i) control and α-actinin overexpressing SC cells for a sharp (A) and a round tip (B), respectively, and (ii) wild-type and vimentin-KO MEFs for a sharp tip (C) and a round tip (D), respectively.

### Results from individual SC cell strains

Fig. S11 shows additional image of filamentous actin (F-actin) and phosphorylated myosin light chain (P-myosin) in confluent SC cells treated with dexamethasone (1μM) as compared with controls Fig. S2 shows the results from individuals SC cells strains treated with dexamethasone and then measured with AFM sharp-tip (A), AFM round-tip (B), TM (C), and OMTC (D). Fig. S3 shows the results from individuals SC cells strains transduced with GFP, α-actinin or RhoA and then measured with AFM sharp-tip (A), AFM round-tip (B), TM (C), and OMTC (D).

### Additional results from finite element modeling

Fig. S4 and S5 show the strain and stress distributions for the same cases considered in Fig. 6, which shows the strain energy distribution. For AFM sharp- tips, strain was localized around sharp tip within cortical network and did not propagate through intracellular network (Fig S4 A, B). The smallest AFM round-tips considered (diameter of 0.8 μm) show the strain propagated into intracellular network and the presence of a stiff cortex (*E_cortex_* = 50* *E_intracellular_*) increased the distance to which strain propagated (Fig S4 C, D). This trend is continued for largest AFM round tip considered (10 μm) with the strain now extending throughout the intracellular network (Fig S4 E, F). The stress fields (Figure S5) show similar trends, but show considerable stress within the cortex in all cases.

For OMTC, the strain propagated through intracellular network but the stiff cortex increased the propagation distance significantly (Fig S4 G-H). In contrast to strain distribution, the stiff cortex lead to a stress distribution localized close to cortical layer in all conditions (Fig S5).

We examined the effect that cell thickness would have on our results. We examined cells with *E_cortex_*=50\**E_intracellular_*, varying cell thickness from 2.75-30 μm. We found that for cells thicker than approximately 5 μm, there was little effect on *E_apparent_* (inset in Fig. 6B) and a minimal effect on strain energy distribution as shown in Fig. S6. Similar studies were done modeling the effect of cell thickness on an AFM 10 μm tip measurements of *E_apparent_* with the results shown in Fig. S9.

One peculiar result in the OMTC studies is that a monotonic relationship was not found between E_*apparent*_ and bead embedding depth (see Fig. 7B). Fig. S7 shows this result in more detail for the case of *E_cortex_*=50\**E_intracellular_*. This result arises because of several competing effects. First, if we consider only the effects of the cortex surrounding the bead and an unbounded cell below, then it would be expected that as the bead is embedded more deeply (thus increasing the radius of the contact area, then E_*apparent*_ should approach *E_intracellular_* (73). However, two effects counter this. First, as the bead is embedded more deeply, the substrate increasingly influences E_*apparent*_ because of the finite cell thickness (see insert in Fig. 7B). Also, the distal cortex (the part of the cortex that does not directly surround the bead) shows an increased concentration of strain energy for deeply embedded beads (Fig. S8). Both of these effects act to increase *E_apparent_* and thereby explain the non-montonic behavior shown in Figs. 6B and S7.

Mijailovic et al. (31) defined a function β that used to determine how the effective shear modulus was affected by embedding depth. Fig. S10 shows a comparison of our calculated values of *β* as compared to the values from Mijailovic et al. (31) for the case when *E_cortex_*= *E_intracellular_* showing good agreement with the differences presumably due to differences in the discretization used to create the finite element mesh. We extended this same definition of β (see methods) for a non-homogenous case, *E_cortex_*=50* *E_intracellular_*, the results of which are also shown in Fig. S10.

### Statistical techniques used

All statistical analyses were done using JMP Pro 13.0.0. Measurements of cell mechanical properties following dexamethasone treatment were fit with “ Fit Model: Standard Least Square” as a function of dexamethasone concentration; the “Center Polynomials” option was not chosen. The results for the three curves fits shown in Figures 2A and 2B are as follows:

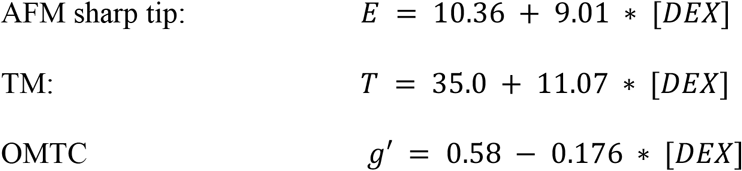

where *E* is modulus (kPa), *T* is RMS traction (Pa), *g′* is the apparent shear modulus (Pa/nm) and *[DEX]* is the dexamethasone concentration μM).

Difference of measurements of cell mechanical properties following induction of GFP, α-actinin, or RhoA were analyzed with “Fit Model: Mixed Model” using the Multiple Comparison option, comparing all pairs with a Tukey correction. In all of these cases, cell strain was included as a random effect attribute to account for within strain variability. Differences of cell mechanical properties between wildtype and knockout MEFS were analyzed with a “t-test” under “Fit Y by X”.

**Fig. S1:**
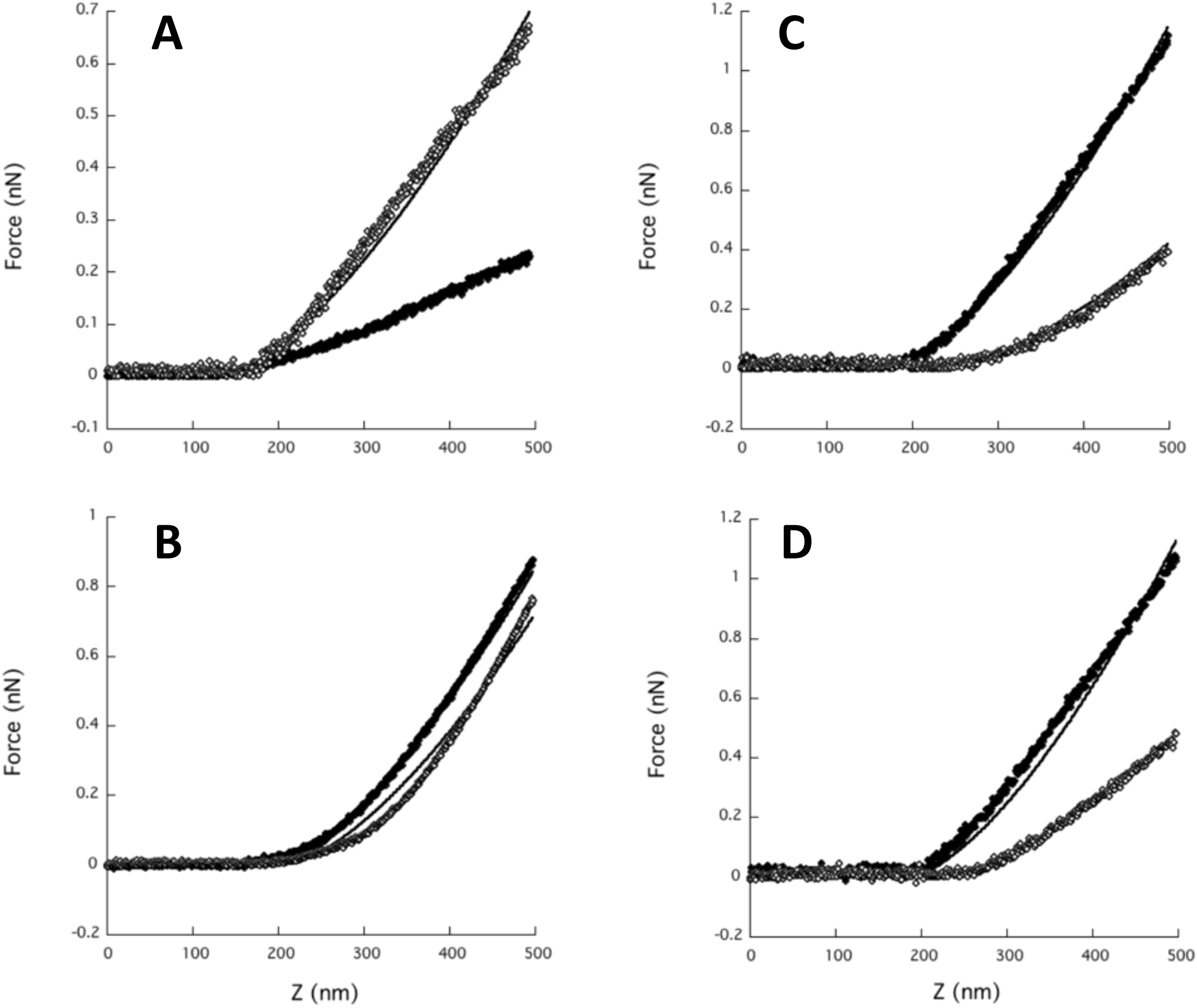
Typical force curves for AFM sharp (panels A, C) and round tips (panels B, D) as a function of ramp distance (Z). Panels A and B compare α-actinin over-expressing SC cells (open symbols) with control SC cells (closed symbols), while Panels C and D compare wild-type MEFs (closed symbols) with vimentin-KO MEFs (open symbols). Fits (solid lines) are from equations (S1:B, D) or (S2: A,C); if line is not visible, it is under the data points.

**Fig. S2:**
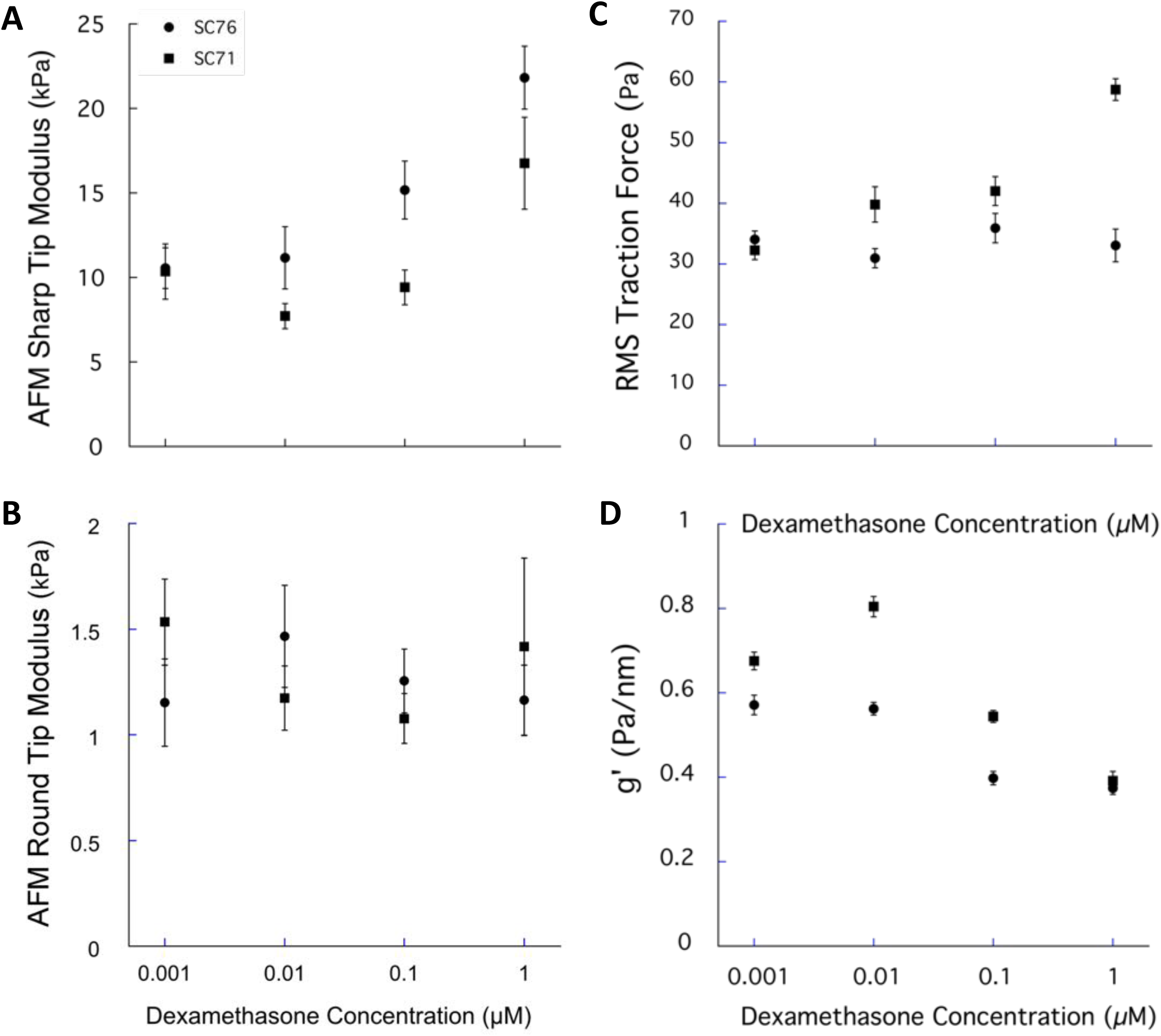
Measurements of cell biophysical properties of confluent SC cells after treatment with varying concentrations of dexamethasone. (A) Sharp AFM tip; (B) Round AFM Tip; (C) TM; (D) OMTC. Geometric Mean ± S.E. about geometric means.

**Fig. S3:**
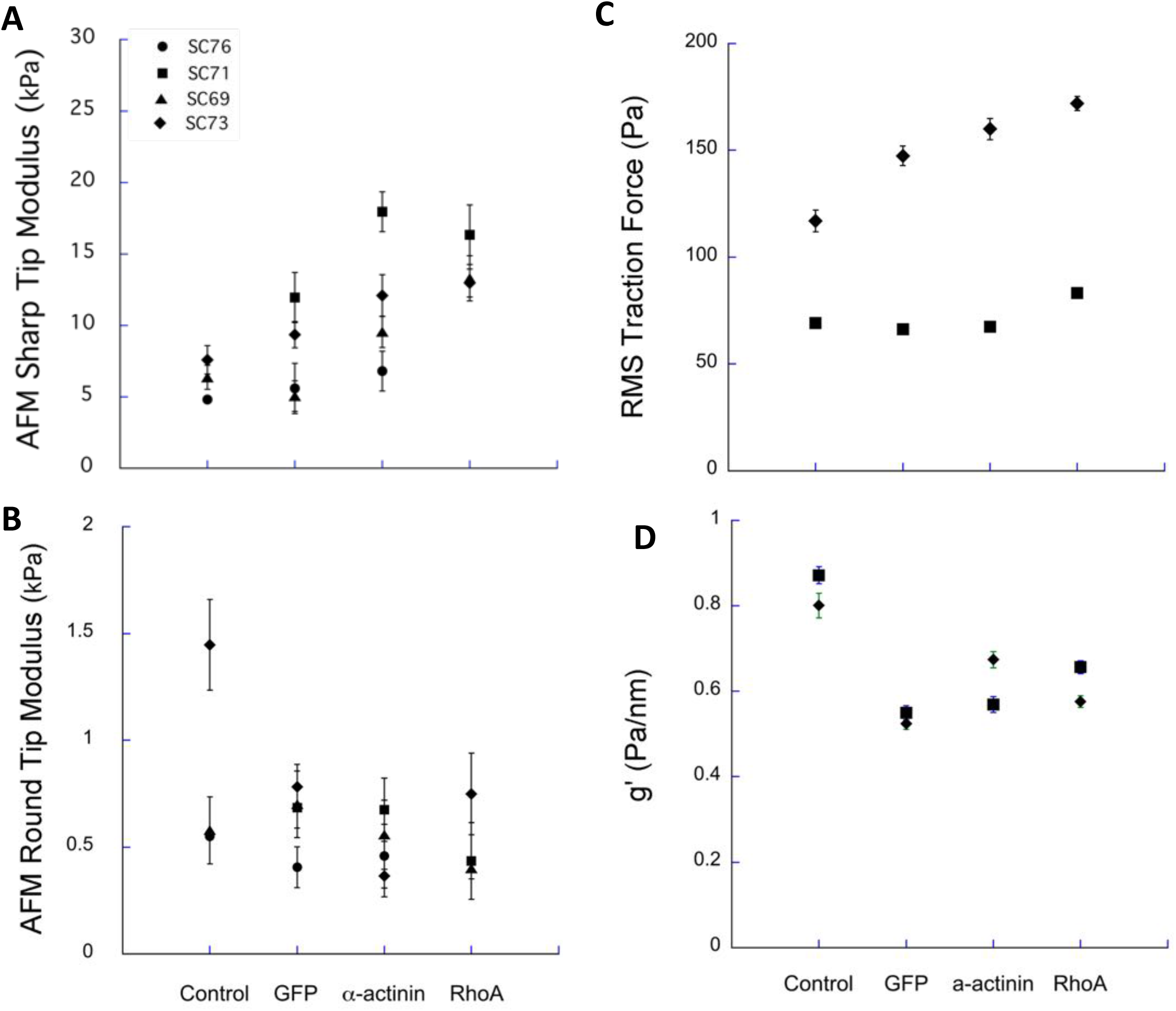
Measurements of cell biophysical properties of subconfluent SC cells after being transduced with GFP, α-actinin or RhoA, as compared with control. (A) Sharp AFM tip; (B) Round AFM Tip; (C) TM; (D) OMTC. Geometric Mean ± S.E. about geometric means.

**Fig. S4:**
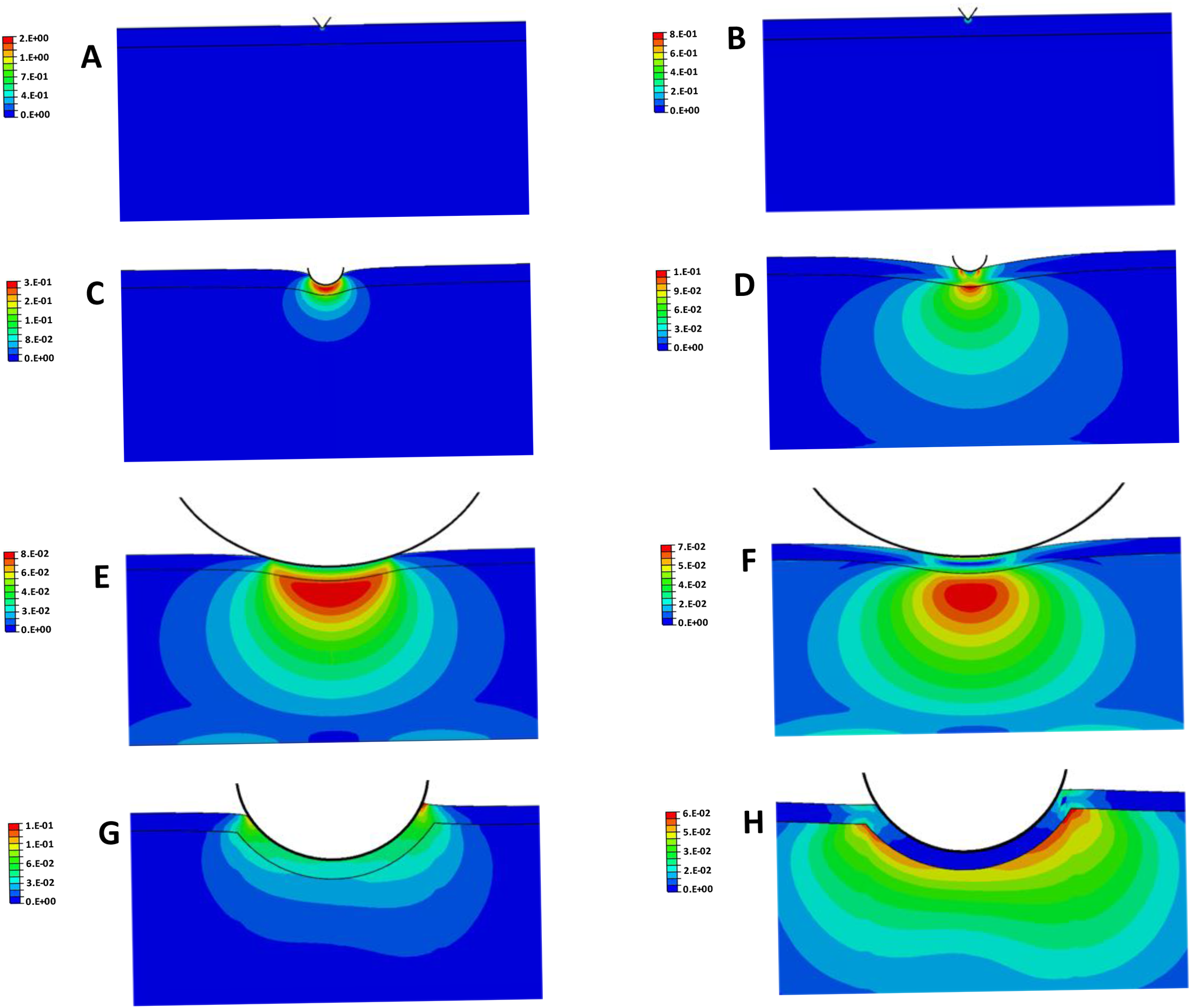
Strain distribution (logarithmic strain) for indentation into a cell of a sharp AFM tip (36° half angle) (A,B), a 0.8 μm diameter rounded AFM tip (C, D), a 10 μm diameter rounded AFM tip (E, F) and a 4.5 μm OMTC bead (G, H). The OMTC bead is embedded 25% of its diameter into the cell and twisted by a torque of 60 Pa applied in a counter-clockwise fashion. Panels A, C, E, G are for cases with *E_cortex_*=*E_intracellular_;* panels B, D, F, H are for *E_cortex_*=50\**E_intracellular_*. Cell thickness is 5 μm. *E_intracelluar_* = 1 kPa for AFM model (12) and 3 kPa for OMTC model (31).

**Fig. S5:**
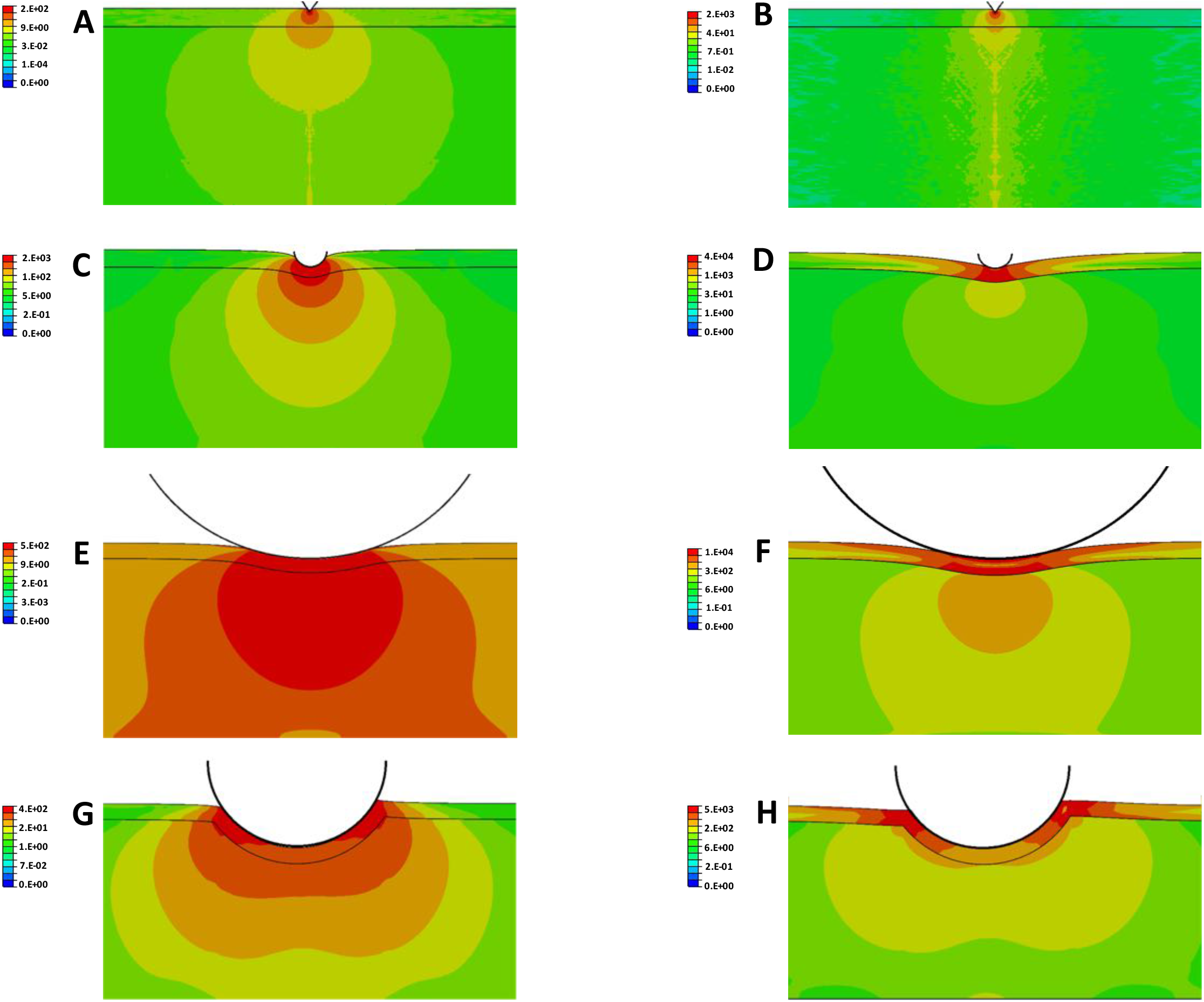
Stress distribution (log scale, Pa) for indentation into a cell of a sharp AFM tip (36° half angle) (A,B), a 0.8 μm diameter rounded AFM tip (C, D), a 10 μm diameter rounded AFM tip (E, F) and a 4.5 μm OMTC bead (G,H). The OMTC bead is embedded 25% of its diameter into the cell and twisted by a torque of 60 Pa applied in a counter-clockwise fashion. Panels A, C, E, G are for cases with *E_cortex_*=*E_intracellular_;* panels B, D, F, H are for *E_cortex_*=50\**E_intracellular_*. Cell thickness is 5 μm. *E_intracelluar_* = 1 kPa for AFM model (12) and 3 kPa for OMTC model (31)

**Fig. S6:**
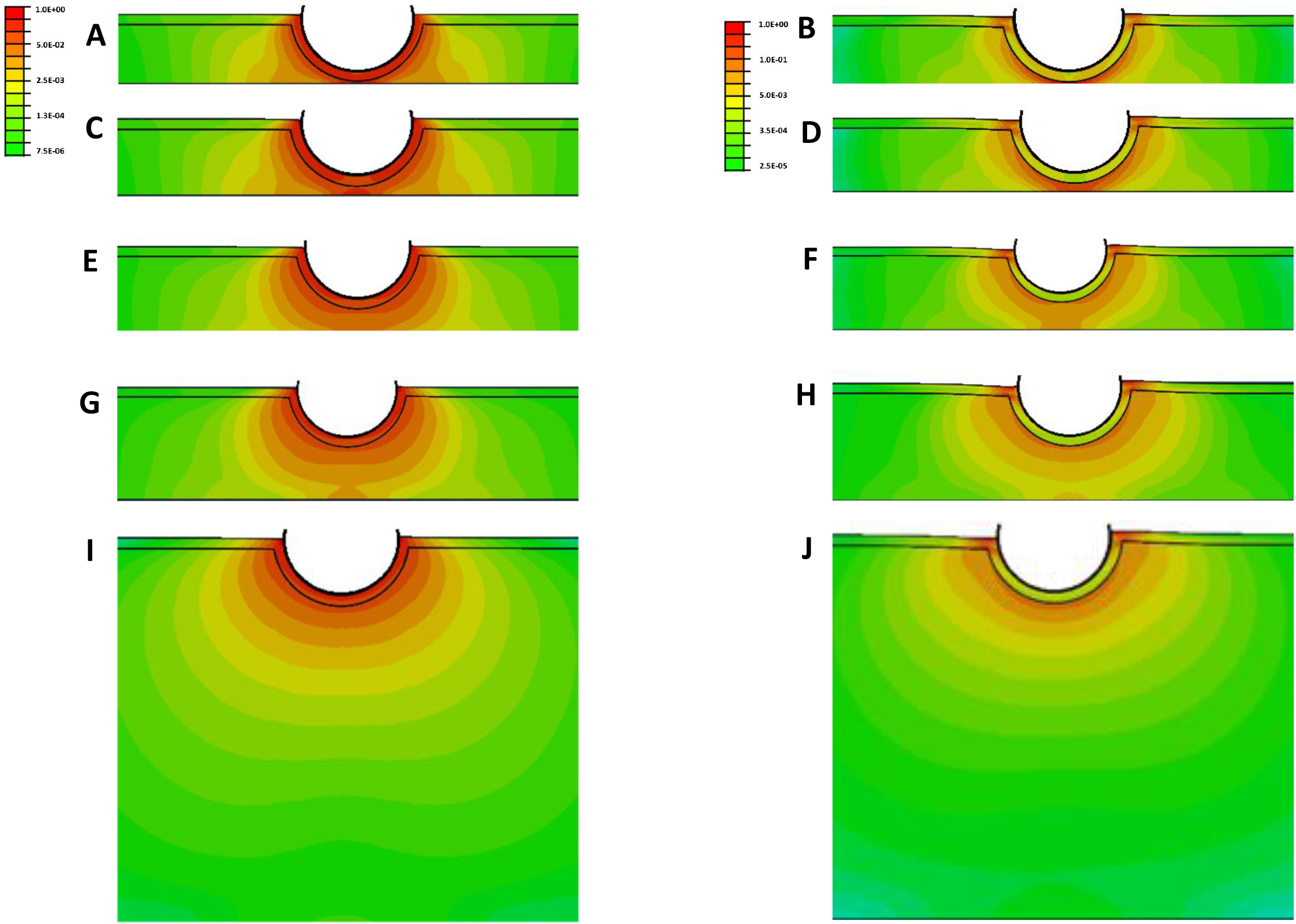
Strain energy density distribution (log scale) for the twisting of a 4.5 μm OMTC bead into cells with *E_cortex_*=*E_intracellular_* (A, C, E, G, I) or *E_cortex_* = 50\**E_intracellular_* (B, D, F, H, J) for cell thicknesses of 2.75 μm (A, B), 3.17 μm (C, D), 4.2 μm (E, F), 5 μm (G, H) and 15 μm (I, J). The strain energy distribution in each panel is normalized to the maximum strain energy in that panel and a log scale is used.

**Fig. S7:**
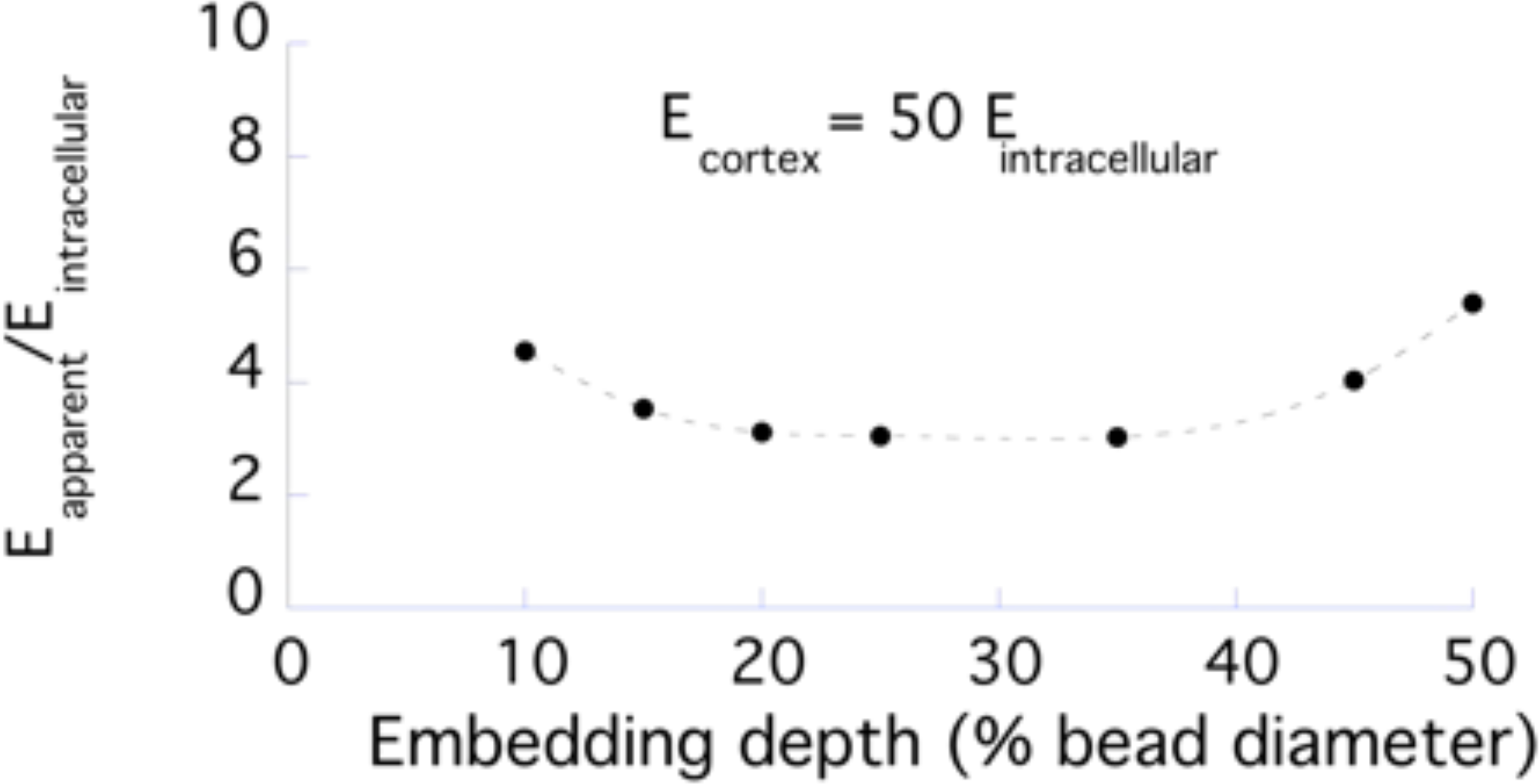
*E_apparent_*/*E_intracellular_* as a function of OMTC bead embedding depth with *E_cortex_*/*E_intracellular_*=50. Cell thickness is 5 μm.

**Fig. S8:**
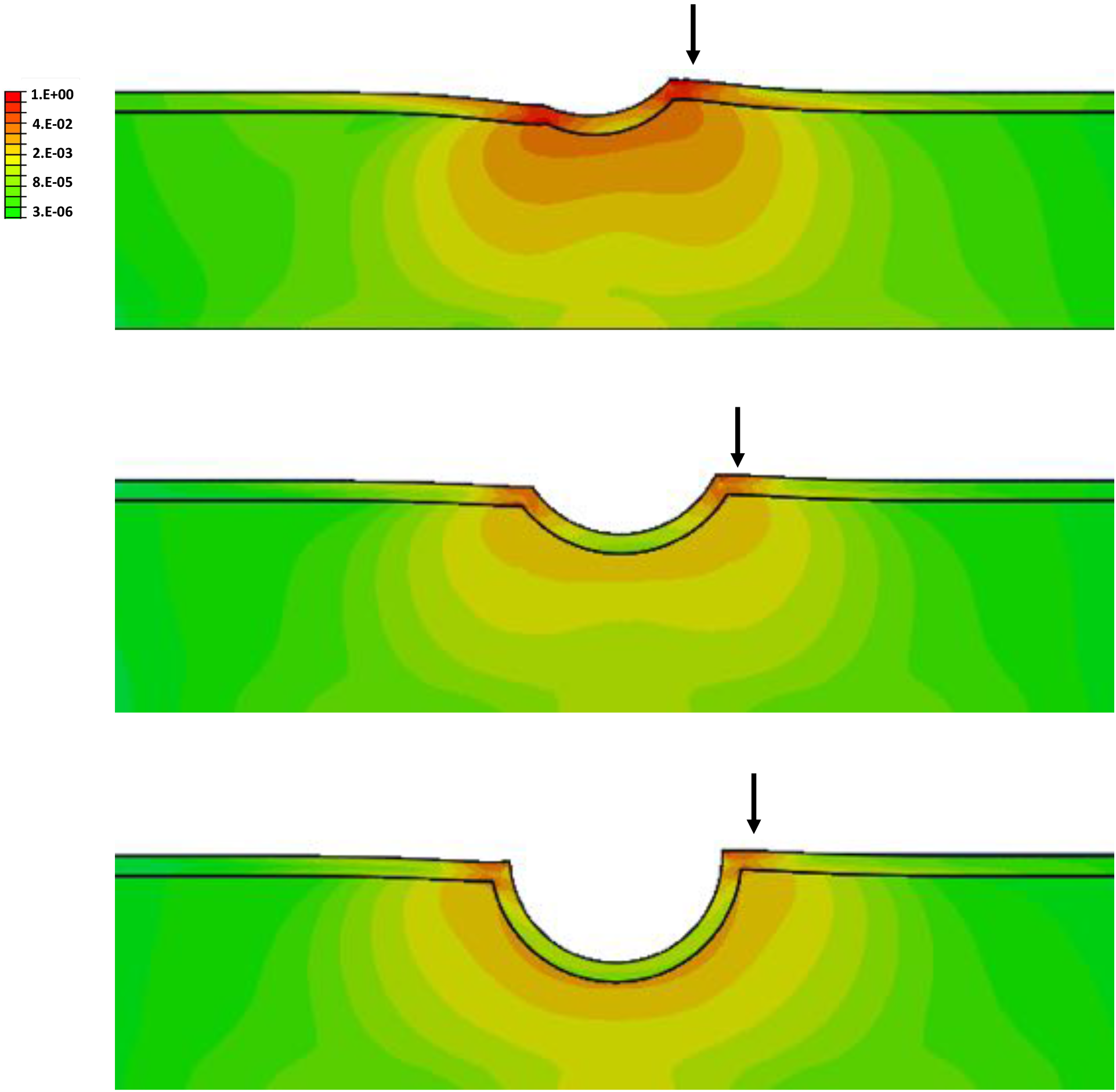
Strain energy distribution (log scale) for the twisting of a 4.5 μm OMTC bead into cells with *E_cortex_*=50\**E_intracellular_* for embedding depths of (A) 10%, (B), 25% and (C) 50%. Arrows point to high strain energy densities in distal cortex. Cell thickness is 5 μm. The strain energy distribution in each panel is normalized to the maximum strain energy in that panel and a log scale is used.

**Fig. S9:**
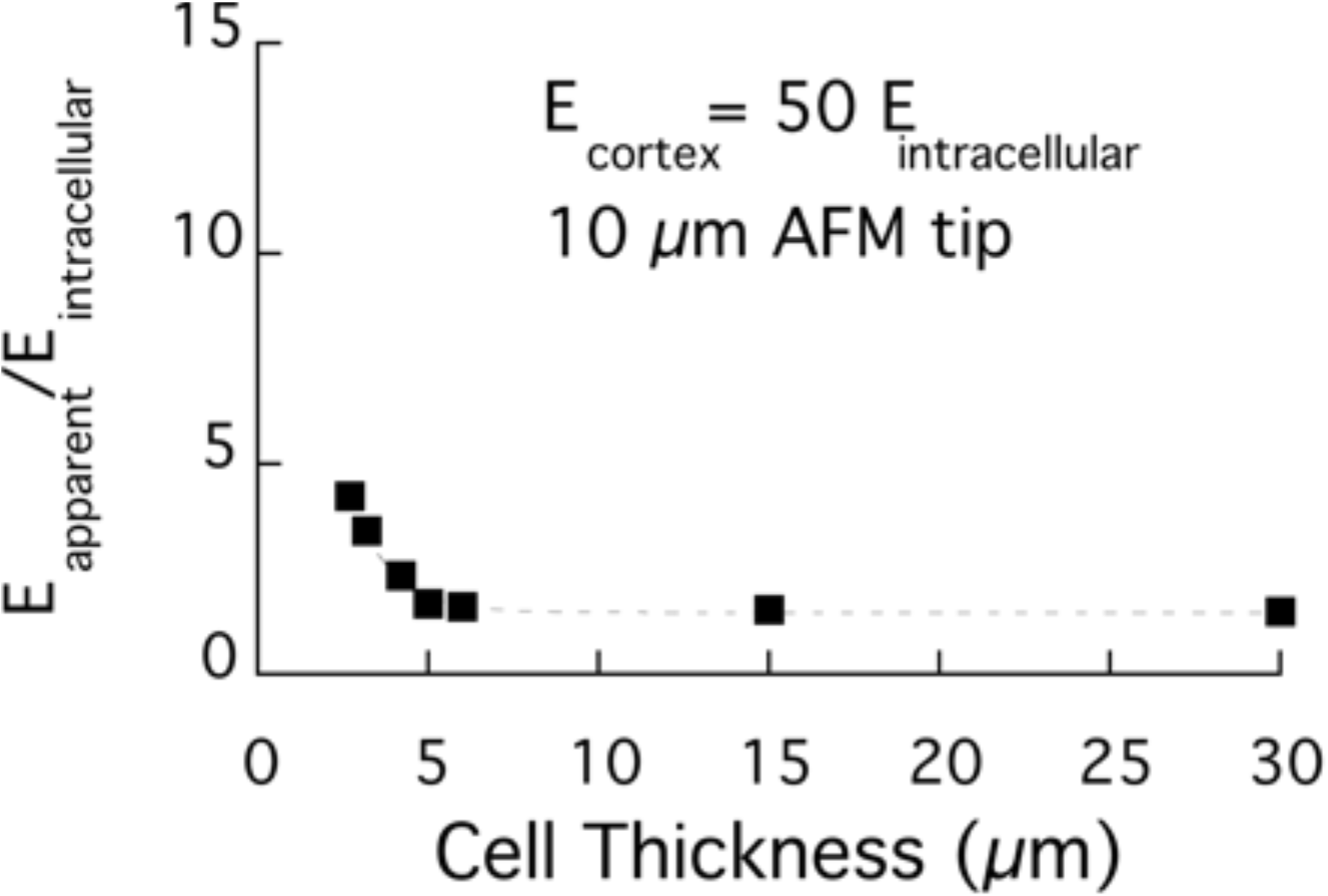
Results for a 10 μm AFM probe indenting 400 nm into a cell with *E_cortex_*/*E_intracellular_*=50 for cell thicknesses ranging from 2.75-30 μm.

**Fig. S10:**
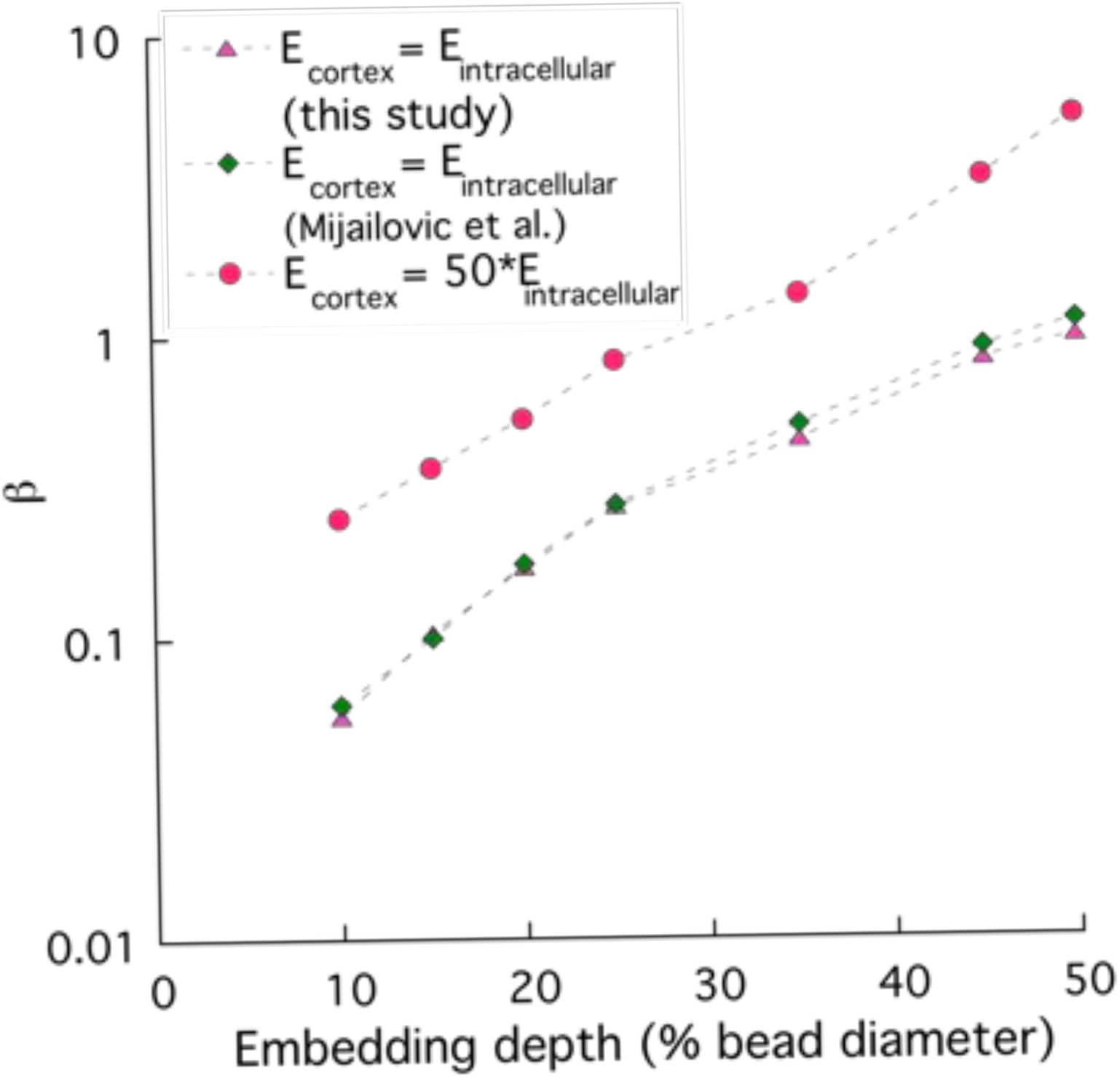
Function β defined by Mijailovic et al. (31) for cell of thickness of 5 μm for case where *E_cortex_* =*E_intracellular_* for this study as compared to the results of Mijailovic et al., and for the case where *E_cortex_* = 50\**E_intracellular_*.

**Fig. S11:**
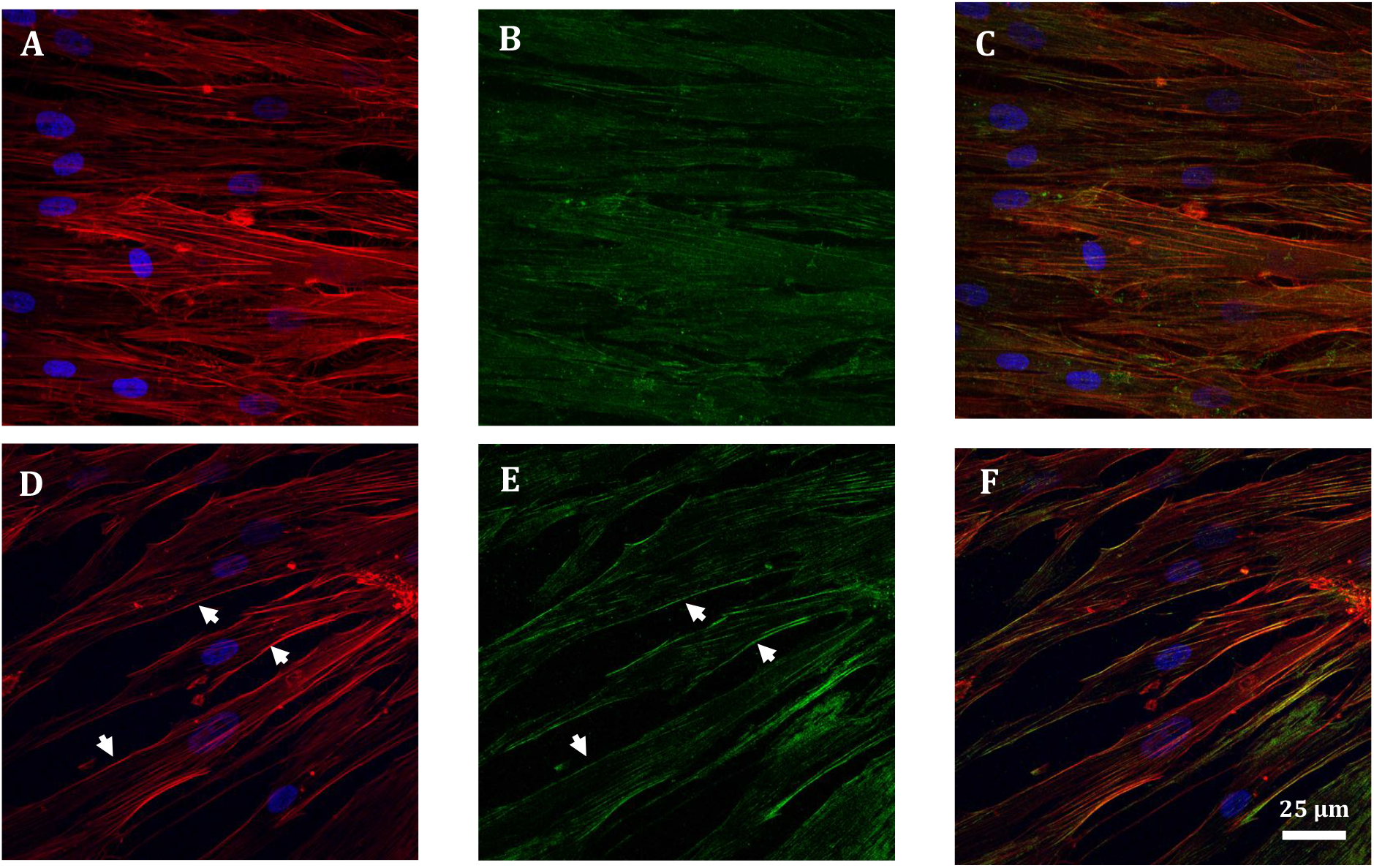
Additional confocal images of confluent monolayer of SC cells treated with vehicle (A-C) and 1μM dexamethasone (D-F); F-actin (red, A & D), phosphorylated myosin (green, B & E), nucleus (blue) and overlaid (C & F). Stress fibers are seen in both groups but the stress fibers are prominent on cortex regions in dexamethasone treated cells (white arrows in D). Also, while p-myosin is present in both groups, it is more concentrated at the cortex of dexamethasone-treated cells (white arrows in E).

